# Patterns of selection reveal shared molecular targets over short and long evolutionary timescales

**DOI:** 10.1101/229419

**Authors:** Jing Li, Ignacio Vázquez-García, Karl Persson, Asier González, Jia-Xing Yue, Benjamin Barré, Michael N. Hall, Anthony D. Long, Jonas Warringer, Ville Mustonen, Gianni Liti

**Author notes:** Correspondence should be addressed to G.L.

## Abstract

Standing and *de novo* genetic variants can both drive adaptation to environmental changes, but their relative contributions and interplay remain poorly understood. Here we investigated the dynamics of drug adaptation in yeast populations with different levels of standing variation by experimental evolution coupled with time-resolved sequencing and phenotyping. We found a doubling of standing variation alone boost the adaptation by 64.1% and 51.5% in hydroxyuea and rapamycin respectively. The causative standing and *de novo* variants were selected on shared targets of *RNR4* in hydroxyurea and *TOR1, TOR2* in rapamycin. The standing and *de novo* TOR variants map to different functional domains and act via distinct mechanisms. Interestingly, standing TOR variants from two domesticated strains exhibited opposite resistance effects, reflecting lineage-specific functional divergence. This study provides a dynamic view on how standing and *de novo* variants interactively drive adaptation and deepens our understanding of clonally evolving diseases.

## Introduction

Darwinian evolution promotes phenotypic adaptation in nature and has important implications in biomedical practices. For example, the emergence of drug resistance during infections and cancer treatment is directly induced by Darwinian evolution in response to drug selection. According to the classic Neo-Darwininstic paradigm, population fitness increases can be attributed to selection favoring beneficial alleles and purging deleterious alleles that are either pre-existing genetic variants segregating in the population before a change in environment (standing variation) or *de novo* mutations emerging after or during an environment change. Beyond scattered examples, the relative contribution to adaptation from these two distinct sources of genetic variation remains poorly characterized (Long, Liti, Luptak, & Tenaillon, 2015).

Connecting allele frequency, phenotype and fitness change in a causally cohesive manner is challenging in both natural and clinical populations, but feasible in experimental populations. Experimental evolution can reveal the molecular determinants of adaptation across a wide range of biological systems with unprecedented resolution (Long et al., 2015). It can be initiated from populations with known levels of standing variation, evolved under fixed selection regimes and preserved *ad infinitum* as frozen fossil records that can be revived and studied in detail. Clonal evolutions of initially isogenic populations have confirmed key theoretical predictions, notably how competing clones carrying different beneficial mutations interfere with each other (clonal interference), and how neutral or slightly deleterious mutations can hitchhike to higher frequencies on the same clone as beneficial mutations (Barrick et al., 2009; Gerrish & Lenski, 1998; Herron & Doebeli, 2013; Kvitek & Sherlock, 2013; Lang et al., 2013; Levy et al., 2015; Payen et al., 2016; Venkataram et al., 2016). Causal relationships in heterogeneous populations, usually derived from sexual crosses of diverged parents, are much more challenging to pinpoint, because of the number of variants that segregate in these populations and the linkage between them. Nevertheless, experimental evolution using heterogeneous budding yeast, fly and Virginia chicken populations have shown that standing variation alone can drive adaptation (Burke et al., 2010; Burke, Liti, & Long, 2014; Parts et al., 2011; Sheng, Pettersson, Honaker, Siegel, & Carlborg, 2015) with no need for *de novo* mutations to emerge and spread. We recently performed experimental evolution using heterogeneous populations derived from diverged West African (WA) and North American (NA) natural yeast strains (hereafter referred to as “two-parent population”), in two different drugs rapamycin (RM) and hydroxyurea (HU) (Vázquez-García et al., 2017). RM is an inhibitor of the eukaryotic serine/threonine kinase TOR and HU is an inhibitor of DNA replication. Specifically, budding yeast contains two TOR genes - *TOR1* and *TOR2.* They form two different complexes termed TOR complex 1 (TORC1) and TORC2. The former contains either *TOR1* or *TOR2* and is uniquely sensitive to RM while the latter specifically contains *TOR2* and is insensitive to RM (Loewith et al., 2002). HU impair DNA synthesis by inhibiting ribonucleotide reductase and preventing the reduction of ribonucleotides to deoxyribonucleotides (Koç, Wheeler, Mathews, & Merrill, 2004). In the two-parent population, we found drug-specific adaptive contributions of standing and *de novo* variants. Selection on standing variation explained more of growth rate increases in HU (51%) than selection on *de novo* mutations (23%) but less in RM (22% vs 70%).

Overall, the relative contribution of standing and *de novo* variants to adaptation depends on multiple factors, including the degree of standing variation, the typical fitness effects of standing and *de novo* variation, the selective constraints imposed by the environment and the relevant time scales (Long et al., 2015). Theory predicts early adaptation in heterogeneous populations to be faster because beneficial standing variants are immediately available and less likely to be lost by drift (Barrett & Schluter, 2008). Standing variants are predicted to disproportionately drive adaptation when *de novo* beneficial mutations are rare, have small selection coefficients, or when the duration of selection is short (Hermisson & Pennings, 2005). However, strict experimental comparisons of adaptation on standing and *de novo* variants are scarce. In particular, it remains to be explored: (1) how the degree of standing variation affects the adaptation rate and yield, (2) whether standing and *de novo* variants are selected in a shared target. These questions have a direct bearing on our understanding of the evolution of resistance to chemotherapy and antimicrobials (Palmer & Kishony, 2013; Turner & Reis-Filho, 2012). To this end, we evolved highly-heterogeneous yeast populations derived from intercrossing four diverged parents over 12 consecutive meiotic generations (Cubillos et al., 2013) (hereafter referred to as “four-parent population”, Figure 1A) to fixed concentrations of RM and HU. In comparison to the two-parent population, the four-parent population has approximately twice the genetic diversity segregating (1 SNP/120bp vs. 1 SNP/230bp), indicating higher level of standing variation (Cubillos et al., 2013). We tracked the adaptation of these four-parent populations to the two drugs at high resolution, comparing the molecular and phenotypic changes to that of the isogenic populations of the four parental strains as well as the published two-parent populations (Vázquez-García et al., 2017). We found that the four-parent populations adapted earlier and faster than the two-parent populations. Resistant standing and *de novo* variants were selected on shared mutational targets (*RNR4, TOR1* and *TOR2).* However, the standing and *de novo* variants of the TOR paralog genes occur in different domains and conferred RM resistance via distinct mechanisms.

**Figure 1.**
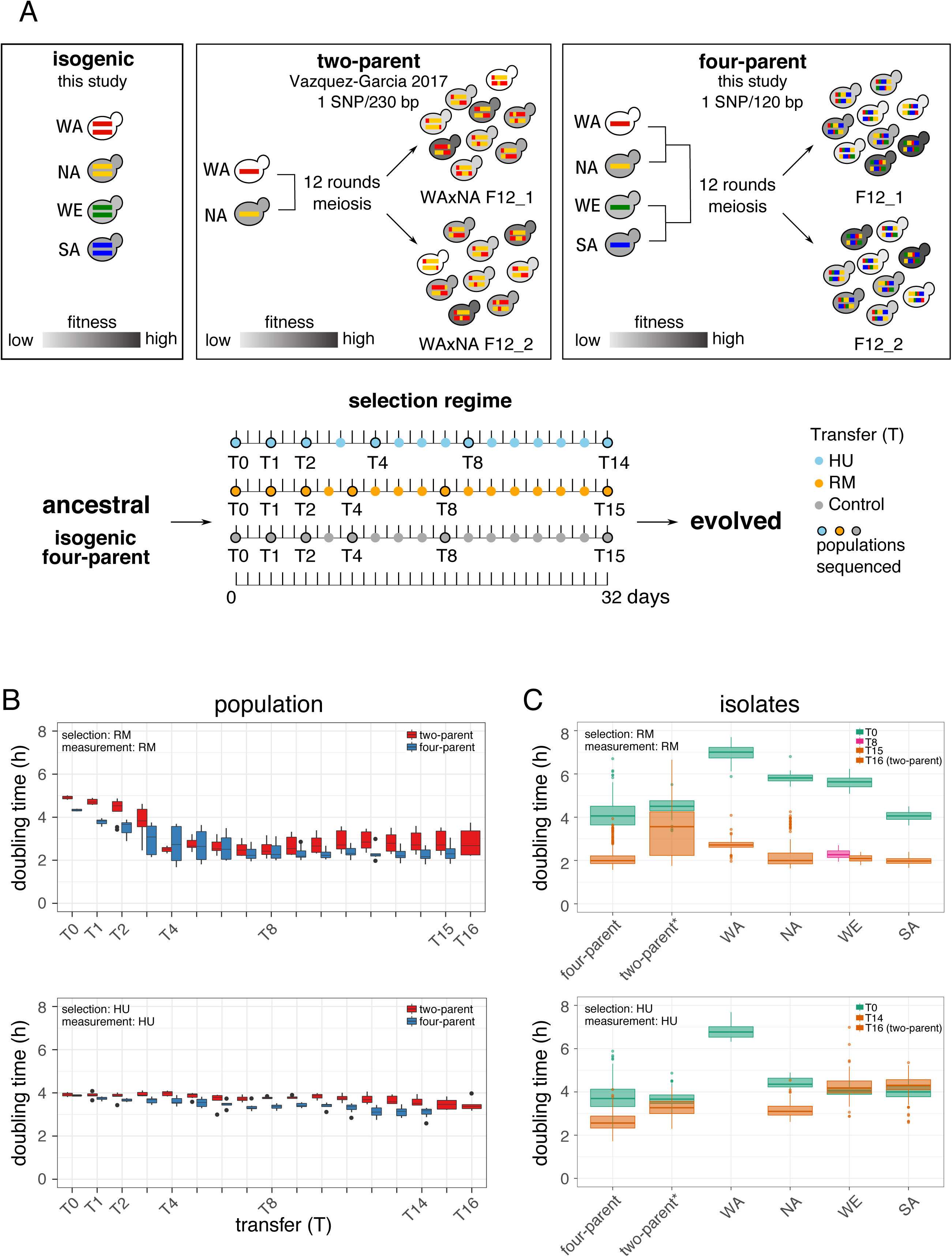
Adaptation of isogenic and heterogeneous populations to rapamycin and hydroxyurea. (A) Ancestral populations with increasing standing variation from isogenic parental, two-parent to four-parent populations (top) and timeline of selection experiment for isogenic and four-parent populations (bottom). The timeline of two-parent selection experiment is listed in Table S1. Random subsamples of the initial populations, and of the 1^st^, 2^nd^, 4^th^, 8^th^ and the last transfer (T14 for HU and T15 for RM in the isogenic and four-parent populations; T16 for HU and RM in the two-parent populations) were sequenced in bulk. (B) Doubling time in RM (top) and HU (bottom) of the randomly sampled bulk populations after each expansion cycle. Boxplot shows the doubling time of all the replicated populations (Table S2). (C) Doubling time of clonal populations expanded from random, single individuals drawn from the ancestral and endpoint populations (Table S2) in RM (top) and HU (bottom). For each drug, we phenotyped 384 random individuals from both the ancestral and endpoint four-parent populations, as well as 48 and 96 random individuals from each ancestral and endpoint isogenic parental replicate population. Boxplot shows the doubling time of these individuals. The WA isogenic populations went extinct after T2 in HU. One WE isogenic population in RM was contaminated at T15 and therefore T8 was analyzed instead. *That wide doubling time distribution of two-parent individuals in RM at T16 is due to the coexistence of fast and slow growth individuals with and without driver mutations, see (Vázquez-García et al., 2017). Boxplot: center lines = median; boxes = interquartile range (IQR); whiskers = 1.5×IQR; points = outliers beyond 1.5×IQR.

## Results

### Adaptation of isogenic and heterogeneous populations to rapamycin and hydroxyurea

To compare adaptation with and without standing variants, we evolved *S. cerevisiae* populations with different levels of standing variation for 32 days (> 50 generations) under RM, HU and basal control condition (no drugs). Four populations (WA, NA, WE, SA – corresponding to strains West African, North American, Wine/European and Sake background respectively) were quasi-homogeneous at the onset of selection, corresponding to clonal expansion of the four diploid parents (Figure 1A, Tables S1-S2). Two four-parent populations (F12_1 and F12_2) were independently derived from the four parents by 12 rounds of intercrossing (Cubillos et al., 2013) and were therefore highly heterogeneous at the onset of selection. We evolved two replicates of each isogenic parental population and eight replicates of four-parent populations in batch-to-batch selection regimes, storing a subsample of each batch (T0 to T14 in HU and T0 to T15 in RM) to create a dense fossil record.

To track the adaptation dynamics comprehensively, we revived the frozen subsamples of all the isogenic, four-parent populations and the previously published two-parent populations (Vázquez-García et al., 2017) across all the time points (Tables S1-S2). We estimated their fitness related properties by both precise measurement of their doubling time and spotting assay (Figures 1B, S1-S3). Over the whole RM experiment, the adaptive gain between four-parent and two-parent populations was similar (45.3% vs. 42.6% of doubling time reduction, Mann–Whitney U-test, *p* = 0.96). However, the early adaptive gain (T0 to T2) was larger in the four-parent populations (19.2% vs. 11.2% of doubling time reduction, Mann–Whitney U-test, *p* = 0.038), highlighting the advantage of higher level of standing variation in driving expeditious adaptation. There was no substantial late stage (last three time points) adaptation in either four-parent or two-parent populations (5.1% of doubling time increase and 1.1% reduction respectively), reflecting exhaustion of adaptive potentials within the experimental timescale. In HU, the adaptation was slow, gradual and persisted to the end in both the four-parent and two-parent populations but with seemingly greater adaptive gains in the four-parent populations (20.4% vs. 12.3% of doubling time reduction, Mann–Whitney U-test, *p* = 0.06). Therefore, a doubling of segregating diversity in the four-parent populations translated into more rapid and more remarkable adaptive gains in both RM and HU. No observable adaptation to control condition (no drug) was observed (Figure S1).

To measure the adaptive gains in individuals independently of their background population, we isolated > 2,600 random clones from ancestral and a subset of the endpoint populations (Table S2) and measured their doubling time separately. Before selection (T0), the variability in doubling time between individuals of the four-parent population was much greater than that in the two-parent populations (Figure 1C, σ^2^ = 0.43 vs. 0.12 in RM and σ^2^ = 0.35 vs. 0.093 in HU). Thus, the higher genetic diversity of the four-parent populations also translated into higher variation in the key fitness component under selection, creating the necessary foundation for faster adaptation. The mean adaptive gain in individuals drawn from four-parent populations at the endpoint also exceeded that of their counterparts from two-parent populations, with a doubling time reduction of 48.2% vs. 27.2% in RM and 29.9% vs. 11.2% in HU. This provides independent verification of the accelerated adaptation in populations with higher level of standing variation (Figures 1C and S4).

Growth phenotyping of both bulk populations (Figures S1 and S3) and individuals drawn from these populations (Figure S4) showed that all initially isogenic populations (NA, SA, WE, WA) achieved certain levels of adaptation to RM. The RM-adapted populations grew faster than their ancestral non-adapted populations regardless of the founding genotype (Figures 1C and S4, Mann–Whitney U-test, *p* < 2.2 × 10^−16^). Individuals drawn from the NA, SA and WE endpoint populations reached the same level of adaptation as those from the four-parent populations, whereas those from the WA populations adapted more slowly, which is consistent with their weaker initial growth. Remarkably, only the NA managed to adapt to HU (28.2% of doubling time reduction, Mann–Whitney U-test, *p* < 2.2 × 10^−16^). Even though NA individuals failed to reach the same adaptation level of four-parent individuals (Figure 1C; mean endpoint doubling time 3.16 vs 2.62 hours, Mann–Whitney U-test, *p* < 2.2 × 10^−16^). The SA and WE individuals grew worse at the end of HU selection than their respective ancestral states (6.5% and 4.2% of doubling time increase). The WA individuals went complete extinction after two cultivation rounds (T2), suggesting the lowest evolvability. In summary, we found that higher level of standing variation positively impacts the rate of adaptation, the absolute adaptive gains and the endpoint performance, with exact effects depending on the selection regime.

### *De novo* mutations in *TOR1* and *FPR1* drive rapamycin adaptation in isogenic populations

To lay a solid foundation for understanding the adaptation of the highly heterogeneous four-parent populations, we sequenced the initially isogenic populations at multiple time points (Tables S1-S3). As expected, *de novo* mutations drove adaptive evolution in isogenic populations. In RM, we detected recurrent mutations in *TOR1* and *FPR1* (Figure 2A). *TOR1* mutations (six mutations in three sites) emerged in all the eight populations, indicating *TOR1* as a background independent source of RM resistance. In contrast, *FPR1* mutations (frame shift and start codon disruption in two sites) emerged only in the two NA populations. Surprisingly, all the NA clones carrying *FPR1* mutations became haploids during selection. This may be a consequence of NA diploids being highly prone to sporulate even in relatively rich medium (Cubillos, Louis, & Liti,2009) and a strong selection for haploids carrying loss-of-function *FPR1* mutations given that they are fully recessive (Vázquez-García et al., 2017). The frequency increase of *TOR1* and *FPR1* mutations agreed well with the doubling time reduction of the populations in which they emerged (Figure 2A). This supports that they are true drivers of adaptation, rather than hitchhikers or drifters, and that adaptation is genetic, rather than initially epigenetic and later genetically assimilated (Gjuvsland et al., 2016).

**Figure 2.**
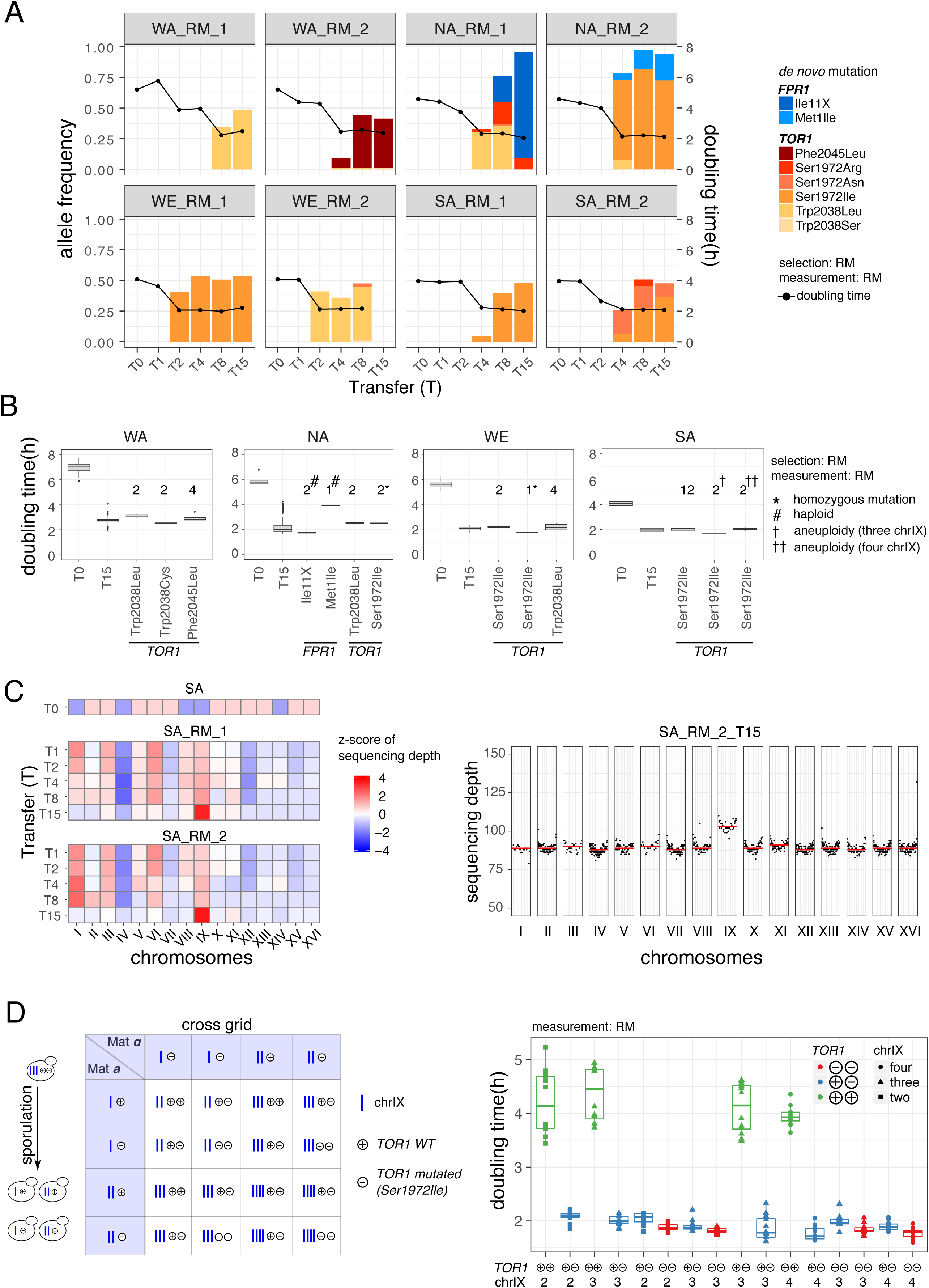
*De novo* mutations in *TOR1* and *FPR1* drive rapamycin adaptation in isogenic populations. (A) Bars: frequency dynamics of *de novo* driver mutations emerging in isogenic populations adapting to RM (left y-axis). Bar color = driver mutations (in *FPR1* = light-dark blue, in *TOR1* = yellow-brown). Line: the mean doubling time of bulk population (right y-axis). (B) Doubling time of random individuals drawn from the ancestral (T0, *n* = 48 for each parent), RM evolved (T15, *n* = 192 for each parent) populations and genotyped individuals. We divided genotyped individuals into groups based on their driver mutations; no individual carried more than one driver mutation. The number above each boxplot indicates the number of genotyped individuals with confirmed driver mutations by Sanger sequencing. (C) Median chromosome sequencing depth (*x*) for each chromosome in isogenic SA populations adapting to RM (left), shown as a *z*-score = (*x* - *μ*)/*σ*, here *μ* and *σ* is the mean and standard deviation of sequencing depth of each population. The genome-wide sequencing depth of population SA_RM_2_T15 (right), measured by whole-population genome sequencing. Genomic positions are shown on the *x*-axis; the sequencing depth is shown on the *y*-axis. Each point indicates the median sequencing depth within a 10-kb window on each chromosome. The red line indicates the median sequencing depth of each chromosome. (D) Design (left) and doubling time (right) of a cross grid experiment. We crossed spores from individuals drawn from the RM evolved (T15) SA populations to generate diploids with known driver mutation genotypes. “+” and “−” = *TOR1* genotypes, WT and *de novo* mutated respectively. Blue bar = chromosome IX. Marker shape = chromosome IX copy number, marker color = *TOR1* genotype. Boxplot: center lines, median; boxes, interquartile range (IQR); whiskers, 1.5×IQR. Data points beyond the whiskers are outliers.

To quantify the individual contributions of *TOR1* and *FPR1* mutations to RM adaptation, we isolated and estimated the doubling time of individual clones carrying these mutations (Figure 2B, Table S4). Except for *FPR1* Met1Ile, the doubling time reduction conferred by each individual mutation in the relevant state (heterozygous or homozygous) equaled (e.g. *TOR1* S1972I in WE), or approached (>90%, e.g. *TOR1* W2038L and S1972I in NA) the total doubling time reduction of the population in which it emerged (Figure 2B). Nearly all the clones from the evolved populations carried one of these mutations; thus, they were capable of explaining almost the complete adaptive gains (Heitman, Movva, & Hall, 1991). All RM-adapted populations performed equally well in presence and absence of RM; thus RM adaptation had plateaued (Figure S4). *TOR1* mutations recurrently emerging in different genetic backgrounds (Ser1972Ile in NA, SA and WE and Trp2038Leu in WA and WE) consistently gave complete tolerance to RM (Figure 2B). The larger adaptive gain conferred by *FPR1* Ile11X frame shift than by *FPR1* Met1Ile stop in the NA background (69.8% vs. 32.9% of doubling time reduction, Mann–Whitney U-test, *p* = 2.7 × 10^−6^, Figure 2B) agreed with the near fixation of *FPR1* Ile11X in NA_RM_1 and the low frequency of *FPR1* Met1Ile in NA_RM_2 (Figure 2A). Given that both should be complete loss-of-function mutations, this distinction is intriguing. In the WE background, the *TOR1* Ser1972Ile homozygous clones grew faster than those with the heterozygous mutation (Figure 2B, 68.1% vs. 59.8% of doubling time reduction, Mann–Whitney U-test, *p* = 1.9 × 10^−4^), giving them a competitive edge and suggesting that continued selection should drive the homozygote state to fixation. Such homozygous mutations should have occurred via loss of heterozygosity, as demonstrated in our previous study (Vázquez-García et al., 2017).

Whole-genome population sequencing uncovered no copy number changes, except in both SA populations where the sequencing depth of chromosome IX (chrIX) increased under RM selection (Figure 2C, Table S3). Copy number qPCR confirmed that the RM-evolved diploid SA clones carried three or four rather than the normal two copies of chrIX. Given that extra chrIX copies conferred dramatically increased heat sensitivity (Figure S5F), we estimated that ∼12.5% and ∼8.3% of the evolved population (SA_RM_2_T15) carried three and four copies chrIX copies respectively based on the frequencies of heat-sensitive clones. This is roughly in agreement with the estimates based on the sequencing depth analysis (Figure 2C). All the SA clones with extra chrIX copies carried the *TOR1* Ser1972Ile heterozygous driver mutation. Mutated *TOR1* clones carrying three copies of chrIX grew faster in RM than those with two or four copies (Mann–Whitney U-test, *p* = 6.90 × 10^−5^ and *p* = 3.43 × 10^−3^ respectively) (Figure 2B). To better understand the interplay between the *TOR1* Ser1972Ile mutation and the chrIX amplification, we constructed a cross grid of diploid strains with all possible combinations of *TOR1* (wild type or mutated) and chrIX copy number (2-4 copies) and estimated their doubling time (Figure 2D). Overwhelmingly, the *TOR1* Ser1972Ile mutation is the major contributor to RM resistance (53.2% and 56.1% of doubling time reduction for heterozygous and homozygous mutation respectively), with extra chrIX copies being marginally beneficial in the *TOR1* Ser1972Ile clones. In sharp contrast to RM selection, isogenic populations propagated under HU almost uniformly failed to generate and maintain detectable *de novo* variants. No *de novo* driver mutations were detected in WE, WA or SA populations, which probably explains the evolution failure of these populations throughout the 32-day experiment (Figures S1, S3-S4). The *RNR4* mutations (Arg34Ile and Lys114Met) were detected in the NA background. The clones carrying these mutations in heterozygous state showed a mean population doubling time reduction of 31.8%.This adaptive gain was directly comparable to those of NA endpoint populations, and thus capable of fully explaining their adaptive gains. (Figure S4B).

### *De novo* mutations in *TOR1, TOR2* and *FPR1* drive rapamycin adaptation in heterogeneous populations

Both *de novo* and standing variants could contribute to adaptation in the four-parent populations. Given that they were derived from four parents, the frequency spectrum of each parental allele is centered around a median frequency of 0.21 (WA), 0.26 (NA), 0.26 (WE) and 0.26 (SA) at T0 (Cubillos et al., 2013). In comparison, the initial frequency of *de novo* mutations is extremely low, arising during the crossing or selection phases (Vázquez-García et al., 2017). We called *de novo* driver mutations in genes that were recurrent mutation targets in the eight four-parent populations and found that *FPR1, TOR1* and *TOR2* harbor such mutations (Figure 3A). Identical *FPR1* mutations occurred in all the replicated populations derived from one intercrossed population F12_1 and the same *TOR1* mutations were found in all the replicated population derived from the other intercrossed population F12_2; thus these drivers emerged during the shared crossing phase and then expanded independently during the selection phase. Validating this assumption, the same haplotype blocks increased in frequency in the replicated populations derived from the same intercrossed population, reflecting expansion of the same clones present at T0 (Figure S6). Similar to the isogenic lines, we found recurrent mutations at the *TOR1* 1972 and 2038 amino acid sites, further confirming that these were the primary RM selection targets and that they arise independently of the genetic context. In one population (F12_2_RM_4) we also found a *TOR2* Ser1975Ile mutation rising to high frequency (Figure 3A). *TOR2* Ser1975 is located in the RM-binding domain and is paralogous to *TOR1* Ser1972, implying that RM driver targets are conserved between *TOR1* and *TOR2* (Helliwell et al., 1994). Isolation and genotyping of single clones from the population containing *TOR2* Ser1975Ile showed that heterozygous and homozygous clones co-exist. This explains its frequency higher than 0.50 in the population. The doubling time of *TOR2* homozygous mutants was significantly shorter than the heterozygous clone in RM (mean: 1.77 vs. 2.18 hours, Mann–Whitney U-test, *p* = 3.9 × 10^−4^). The doubling time of all the other genotyped mutants, including *TOR1* Ser1972Asn (1.95 hours), Ser1972Arg (2.34 hours), Trp2038Ser (1.93 hours) and Trp2038Leu (1.83 hours) clones in their heterozygote states and *FPR1* Thr82Pro homozygotes (2.18 hours) was faster than that of the clones drawn from the adapted populations but not carrying these driver mutations (3.45 hours), suggesting clear phenotypic contributions from these *de novo* driver mutations.

**Figure 3.**
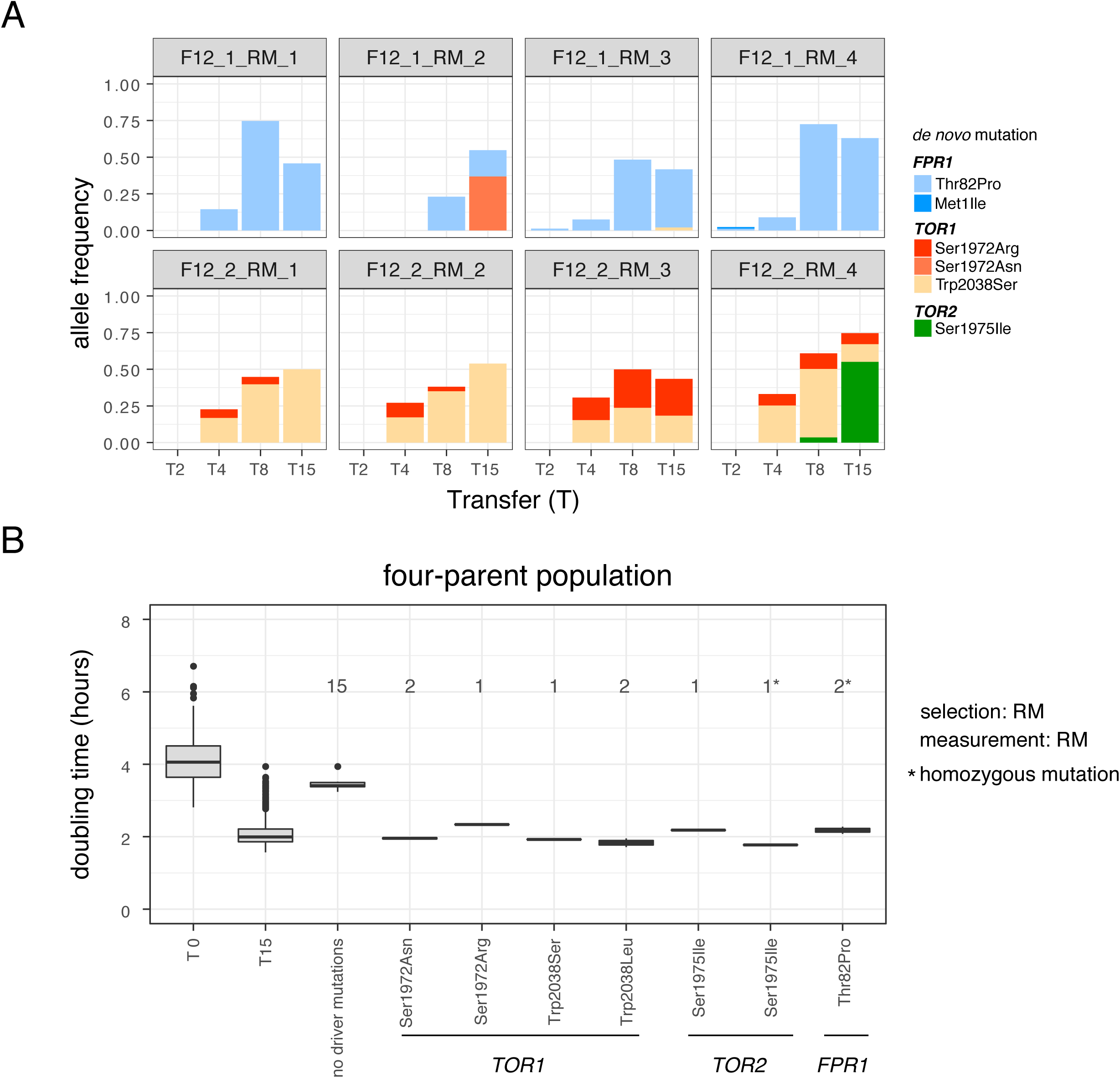
*De novo* mutations in *TOR1, TOR2* and *FPR1* drive rapamycin adaptation in heterogeneous populations. (A) Frequency dynamics of *de novo* driver mutations emerging in four-parent populations adapting to RM. Top and bottom panels show replicates from F12_1 and F12_2 respectively. (B) Doubling time of random individuals drawn from the ancestral (T0, *n* = 384) and RM evolved (T15, *n* = 384 individuals) populations. We divided genotyped individuals into groups based on their driver mutations; no individual carried more than one driver mutation. The number above each boxplot indicates the number of genotyped individuals with or without driver mutations by Sanger sequencing. Boxplot: center lines, median; boxes, interquartile range (IQR); whiskers, 1.5×IQR. Data points beyond the whiskers are outliers.

In stark contrast to RM, we did not detect any *de novo* driver mutations in the four-parent populations under HU selection. Nevertheless, adaptation to HU is obvious (Figures 1B-C, S4D), and genome-wide frequency changes of parental alleles showed broad jumps at later time points (Figure S7), indicating resistant clones rising to high frequencies. We therefore conjectured that standing variation largely drove the adaptation to HU in the four-parent populations.

### Standing variation provides multiple selection targets to drive adaptation in heterogeneous populations

We next investigated how the standing variation in the four-parent populations contributed to RM and HU adaptation. We searched for genomic regions (quantitative trait loci, QTLs) with a consistent change in the frequency of one or more alleles across both time points and replicated populations. At later time points (T4 to T15), we observed strong shifts of allele frequencies over large genomic regions, reflecting drug-resistant clones rising to high frequency in both selection regimes (Figures S7-S8). Therefore, we analyzed allele frequency changes before the clones arose (T0-T4 for HU and T0-T2 for RM) to map QTLs using 99% and 95% quantiles cut-offs (see Materials and Methods).

In HU, two QTLs passed the 99% quantile cut-off and seven more QTLs passed the 95% cut-off (Figure 4A and Table 1) with a median size of 22 kb and containing on average 10 genes. The peak of one strongest QTL (chrVII: 841~863 kb) coincided with the location of the *RNR4* gene, encoding the small subunit of ribonucleotide-diphosphate reductase that is inhibited by HU. The *RNR4^WE^* allele was selected over the other three parental alleles throughout the selection experiment (Figure 4B-C). We experimentally validated the selective advantage of the *RNR4^WE^* allele by reciprocal hemizygosity (Warringer, Liti, & Blomberg, 2017), finding it to account for an 11.7% of doubling time reduction in the NA/WE diploid background in HU (Figures 4D, S5A-B). This corresponds to ~50% the total doubling time reduction in HU-adapted four-parent populations. The other strong QTL (chrIV: 503∼563 kb) encompassed the highly pleiotropic *ENA1, ENA2,* and *ENA5* transporter genes (Warringer et al., 2011) with the SA allele driving towards fixation in all replicate populations (Figure S9). Four of the seven QTLs passing the 95% quantile showed continuous allele frequency changes until the end of the selection (Table S5, Figure S9) while the allele frequency changes of the other three QTLs wore off before the end of selection. Given that there were no detectable *de novo* driver mutations, the latter was probably due to overwhelming competition from clones carrying the beneficial versions of the stronger QTLs (e.g. *RNR4^WE^, ENA^SA^).*

**Figure 4.**
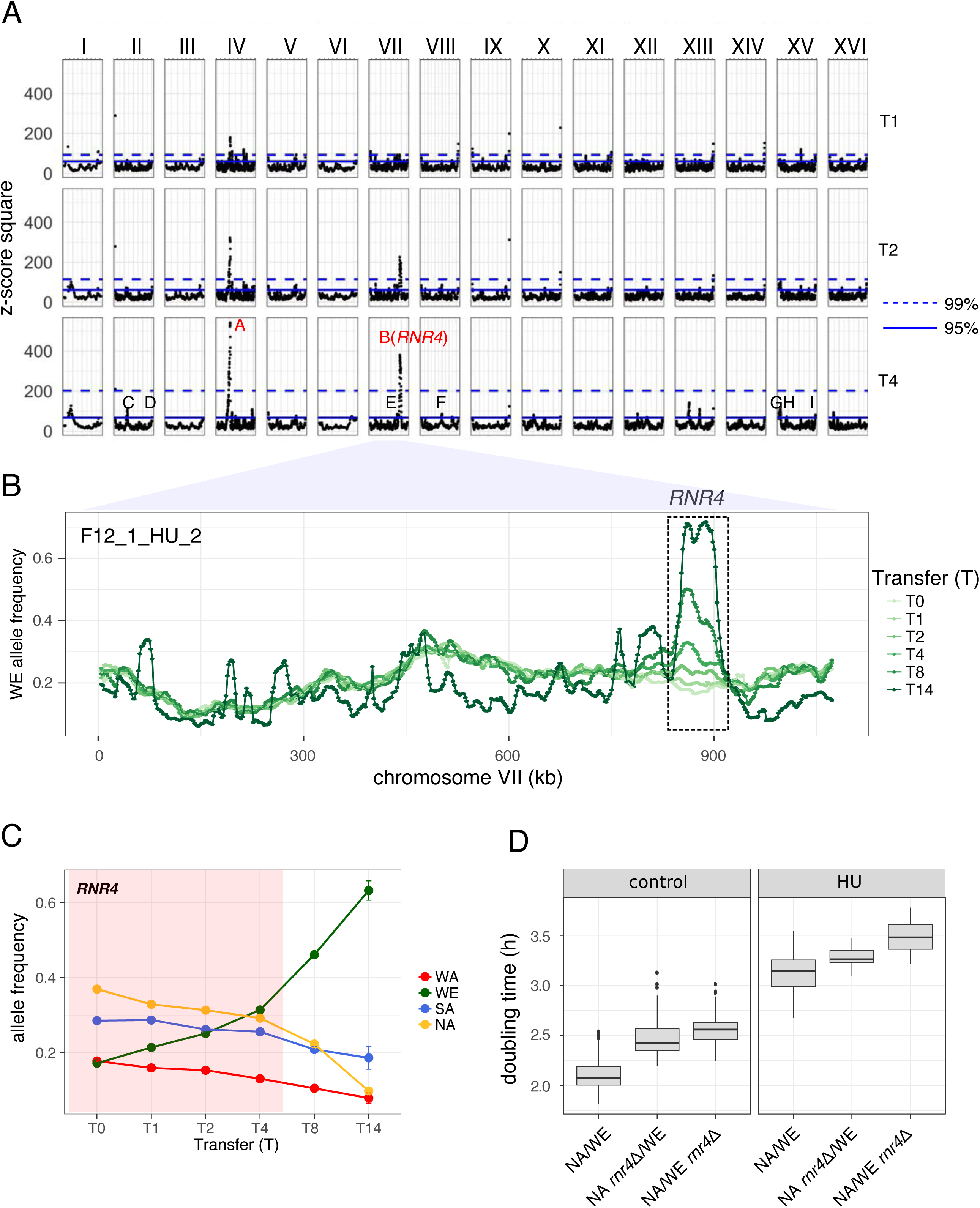
*RNR4* QTL drive adaptation in heterogeneous populations in HU. (A) The *z*-score square is derived from allele frequency changes compared to T0 during early phase of selection (T1-T4) and underlies QTLs. Dashed and solid lines indicate 99% and 95% quantile cut-off respectively. Strong QTLs are labeled in red and weak ones are in black (coordinates listed in Table 1). (B) WE allele frequency changes in chromosome VII in one of the four-parent populations evolved in HU (F12_1_HU_2) from T0 to T14. The region in the black box contains the *RNR4* QTL. (C) Frequency changes of the four *RNR4* alleles from T0 to T14, showing 1:3 segregating pattern (one strong allele vs. three weak alleles). The error bars indicate the standard deviation of all the eight replicates. The region highlighted in red indicates the early phase of selection used for QTL mapping. (D) Doubling time of *RNR4* reciprocal hemizygotes measured in HU and control experimentally confirmed the *RNR4* causative variants. Boxplot: Center lines, median; boxes, interquartile range (IQR); whiskers, 1.5×IQR. Data points beyond the whiskers are outliers.

**Table 1.**
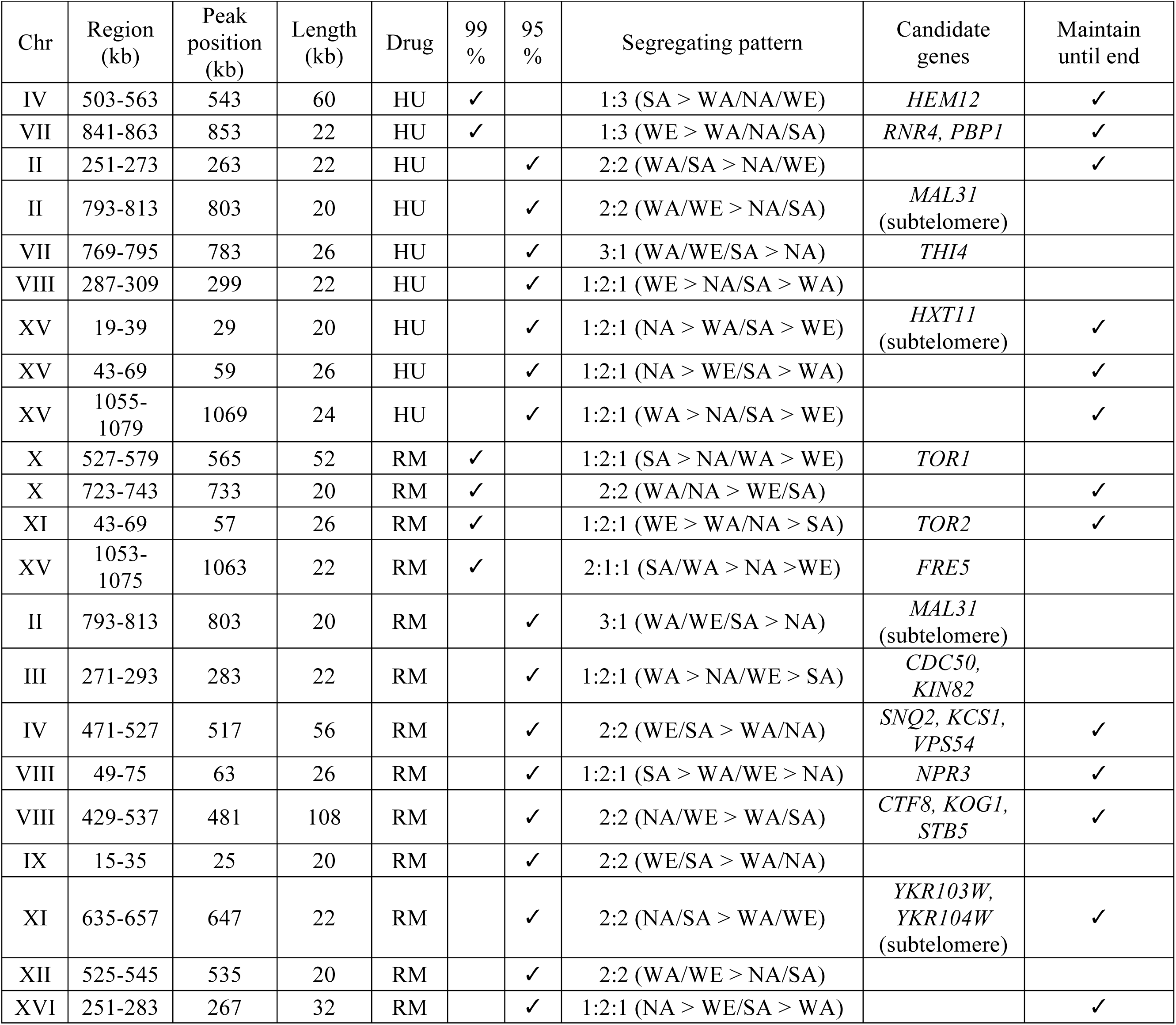
List of quantitative trait loci (QTLs)

Similarly, we mapped QTLs for RM resistance by analyzing allele frequency changes from T0 to T2. We identified four QTLs at the 99% quantile cut-off (Table 1, Figure S10A). The two strongest QTLs (52 and 26 kb respectively) covered the *TOR1* and *TOR2* genes respectively. Interestingly, the WE and SA alleles of *TOR1* and *TOR2* showed opposite allele frequency changes: *TOR1^SA^* and *TOR2^WE^* were selected for while *TOR1^WE^* and *TOR2^SA^* were selected against (Figure 5A). We validated such parental-specific allele preference by reciprocal hemizygosity (Figures 5B, S5C-D). Clones carrying the strong *TOR1^SA^* showed significantly shorter doubling time and higher yield than the ones with the weak *TOR1^WE^* allele (Mann–Whitney U-test, *p* = 3.1 × 10^−4^ and *p* = 1.5 × 10^−4^ respectively). Clones carrying strong *TOR2^WE^* allele showed significantly higher yield than clones carrying *TOR2^SA^* (Mann–Whitney U-test, *p* = 1.5 × 10^−4^). Nine additional QTLs passed the 95% quantile cut-off. While we have not experimentally validated their effects, we considered *SNQ2, NPR3, KOG1* and *CFT8* to be strong candidates for driving these QTLs based on previous studies. Among them, *CFT8* also contributed to RM resistance in the two-parent populations (Vázquez-García et al., 2017). *SNQ2* encodes a multi-drug resistance ABC transporter, and *NPR3* and *KOG1* act together with TOR in nutrient signaling. Several other QTLs were in subtelomeric regions, with the one at chrXI-R containing the subtelomeric genes *YKR103W* and *YKR104W* that encode multi-drug resistance-associated proteins (Mason, Mallampalli, Huyer, & Michaelis, 2003). Based on the end-to-end genome assemblies of the four parental strains (Yue et al., 2017), we found that these two subtelomeric genes were absent in the WE subtelomere, potentially explaining its dramatic allele frequency decrease. The strong RM QTLs such as *TOR1, TOR2, NPR3, CTF8* and *SNQ2* persisted until late time point in RM (Tables 1 and S5, Figure S9), despite the frequency increase of clones carrying *de novo* driver mutations.

**Figure 5.**
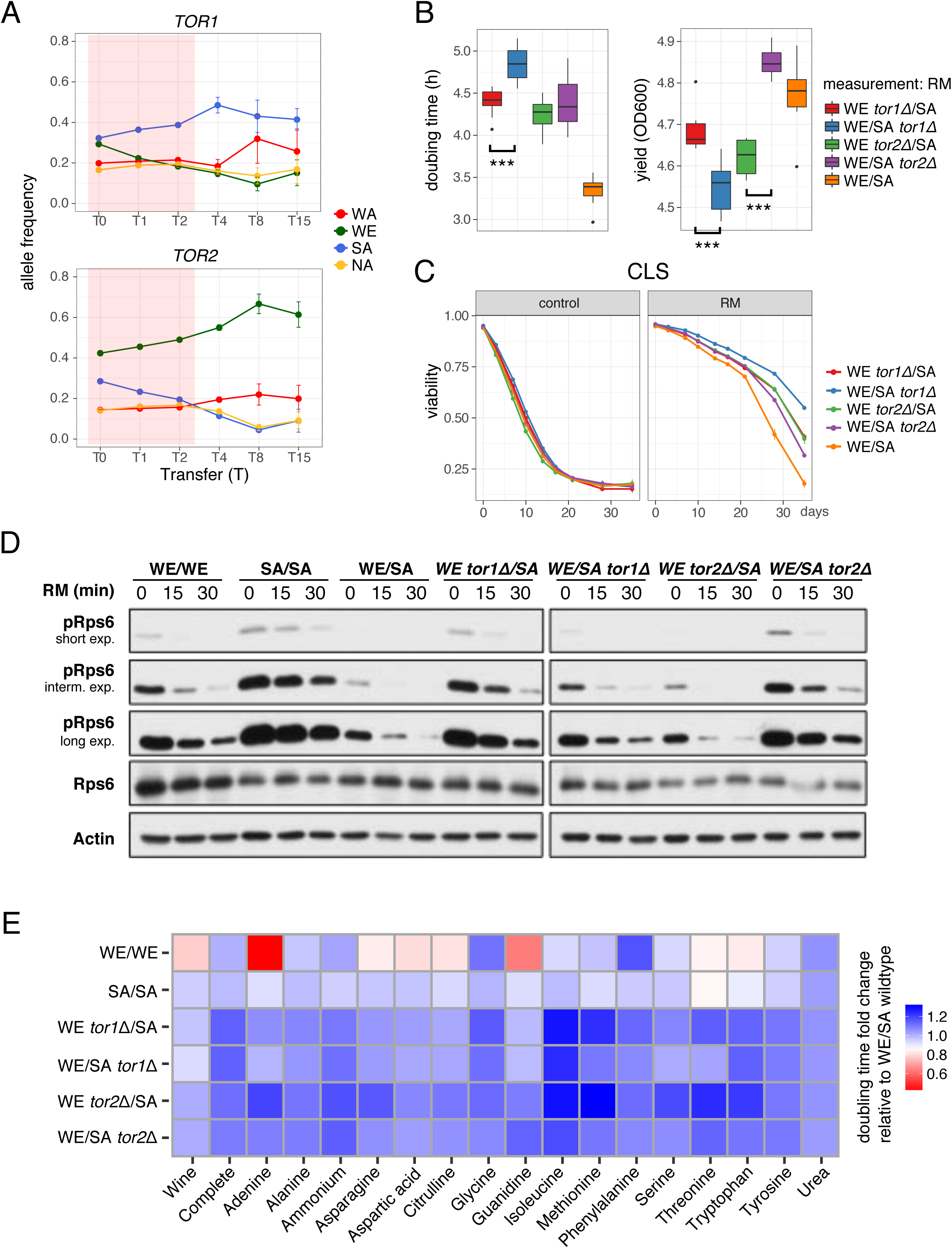
*TOR1* and *TOR2* allelic variation. (A) *TOR1* (top) and *TOR2* (bottom) allele frequency changes of the four-parent populations during RM selection. The region highlighted in red indicates the early phase of selection used for QTL mapping. The points and error bars indicate the mean and standard deviation of all the eight replicates. (B) Doubling time (left) and yield (right) of WE/SA hybrid with *TOR1* and *TOR2* reciprocal hemizygote deletions confirm the causative variants for RM resistance. (C) Chronological life span (CLS) of *TOR1* and *TOR2* reciprocal hemizygotes (WE/SA) in the presence and absence of RM. (D) Characterization of the TORC1 activity by immunoblot of Rps6 phosphorylation in WT parents, hybrid and *TOR1, TOR2* reciprocal hemizygotes. Cells were treated with RM (200 ng/ml) for the indicated time (minute).Total lysates were resolved by SDS-PAGE on 10% polyacrylamide gels and analyzed by immunoblot. Actin was used as loading control. The “short”, “interm.” and “long” panels indicate the exposure time of the membrane to the film. (E) Growth phenotypes of wild type strains and *TOR1, TOR2* reciprocal hemizygotes in 18 environments, corresponding to synthetic wine must and single nitrogen source environments at nitrogen limiting concentrations. Heat map shows the fold change of doubling time compared with WE/SA wild type hybrid.

### Shared selection targets between standing and *de novo* variants in *RNR4, TOR1* and *TOR2*

The multi-hit *de novo* mutations and QTLs identified in the same genes (*RNR4* in HU and *TOR1, TOR2* in RM) showed a pattern of selection on shared molecular targets over short and long evolutionary timescales. To understand why this pattern arose, we compared the standing variants with *de novo* mutations identified in isogenic, two-parent and four-parent populations (Table 2). The HU-resistant *RNR4^WE^* allele had a single derived amino acid change, Ala161Thr, located within the ribonucleotide reductase domain; this substitution was predicted to be functional critical by sequence conservation analysis (see Materials and Methods). The *RNR4 de novo* driver mutations emerging in both NA and two-parent populations were in the same domain but the exact sites differed (Arg34Ile, Arg34Gly, and Lys114Met). All the *de novo* mutations in the *TOR1, TOR2* paralogs occurred in the highly conserved RM-binding domain, where they prevented the binding of the FKBP12-RM complex and thereby conferred RM resistance. In sharp contrast, none of the *TOR1* and *TOR2* standing variants mapped in the RM-binding domain and occurred outside any characterized functional domains with the exception of *TOR1^WE^* Phe1640 (Table 2). Three derived amino acid changes were unique to the weak *TOR2^SA^* allele (Glu122Gly, Ile1369Met, Ile1872Leu) and were all predicted to be deleterious (Table 2). To expand our understanding of natural genetic variation of these shared selection targets, we compared the sequences of *RNR4, TOR1* and *TOR2* across >1,000 *S. cerevisiae* natural isolates (http://1002genomes.u-strasbg.fr/). All the three genes were well conserved (Figure S11). A total of 9, 79 and 73 amino acid sites of *RNR4, TOR1* and *TOR2* respectively were predicted to be functionally critical, based on sequence conservation (Table S6). All nine *RNR4* sites were in the ribonucleotide reductase domain in which the standing and *de novo* variants driving HU adaptation were located. About 38.0% (30/79) of the amino acid sites in *TOR1* and 46.6% (34/73) in *TOR2* were located in known domains, including four *TOR1* and two *TOR2* sites in the RM-binding domain. We experimentally confirmed that natural alleles *TOR1* His2000 and *TOR2* Leu2047 in the RM-binding domain conferred RM resistance (Figure S5E). This adds additional support to that drug resistance can emerge through selection on existing natural variants that prevent drug binding, with no need for *de novo* mutations to emerge.

**Table 2.** Predicting mechanistic consequences of substitutions in genes of interest.

| Gene | Position | Standing variation |  |  |  |  |  | *De novo* mutations | Domain | Score (x10<sup>-2</sup>) | IC |
| --- | --- | --- | --- | --- | --- | --- | --- | --- | --- | --- | --- |
|  |  | *S. par* | S288C | WA | WE | SA | NA |  |  |  |  |
| *RNR4* | 24 | D | D | D | **N** | D | D |  |  |  |  |
| *RNR4* | 161 | A | A | A | **T** | A | A |  | Ribonucleotide reductase small | 1.87 | 2.78 |
| *RNR4* | 114 | K |  |  |  |  |  | M (NA) | Ribonucleotide reductase small | 0.15 | 2.80 |
| *RNR4* | 34 | R |  |  |  |  |  | I (NA, two-parent) | Ribonucleotide reductase small | 0.12 | 2.71 |
| *RNR4* | 34 | R |  |  |  |  |  | G (two-parent) | Ribonucleotide reductase small | 1.70 | 2.71 |
| *TOR1* | 58 | G | D | G | **D** | G | G |  |  |  |  |
| *TOR1* | 133 | S | S | N | **S** | N | N |  |  |  |  |
| *TOR1* | 175 | I | V | **L** | V | V | V |  |  |  |  |
| *TOR1* | 1117 | S | S | S | S | **P** | S |  |  |  |  |
| *TOR1* | 1292 | E | G | G | **E** | G | G |  |  |  |  |
| *TOR1* | 1451 | V | V | **I** | V | V | V |  |  |  |  |
| *TOR1* | 1640 | A | F | V | **F** | V | V |  | FAT |  |  |
| *TOR1* | 1868 | K | K | K | K | **R** | K |  |  |  |  |
| *TOR1* | 1972 | S |  |  |  |  |  | R (NA, SA, four-parent) | rapamycin binding | 4.14 | 2.74 |
| *TOR1* | 1972 | S |  |  |  |  |  | N (SA, four-parent) | rapamycin binding | 0.64 | 2.74 |
| *TOR1* | 1972 | S |  |  |  |  |  | I (WE, SA, NA, two-parent) | rapamycin binding | 0.16 | 2.74 |
| *TOR1* | 2038 | W |  |  |  |  |  | L (WA, WE, NA, two-parent) | rapamycin binding | 0.00 | 2.74 |
| *TOR1* | 2038 | W |  |  |  |  |  | S (four-parent) | rapamycin binding | 0.00 | 2.74 |
| *TOR1* | 2045 | F |  |  |  |  |  | L (WA) | rapamycin binding |  |  |
| *TOR1* | 2091 | A | A | A | A | A | **V** |  |  |  |  |
| *TOR1* | 2414 | R | K | R | **K** | R | R |  |  |  |  |
| *TOR2* | 38 | H | H | **N** | H | H | H |  |  |  |  |
| *TOR2* | 122 | E | E | E | E | **G** | E |  |  | 1.84 | 2.76 |
| *TOR2* | 379 | A | A | A | **S** | A | A |  |  |  |  |
| *TOR2* | 607 | P | S | P | P | **S** | P |  |  |  |  |
| *TOR2* | 1369 | I | I | I | I | **M** | I |  |  | 1.02 | 2.75 |
| *TOR2* | 1975 | S |  |  |  |  |  | I (four-parent) | rapamycin binding | 0.17 | 2.75 |
| *TOR2* | 1856 | I | I | I | I | **T** | I |  |  |  |  |
| *TOR2* | 1872 | I | I | I | I | **L** | I |  |  | 2.16 | 2.75 |

3 We aligned the sequences of *RNR4*, *TOR1* and *TOR2* from the four parents, the *S. cerevisiae* 4 reference strain S288C and *S. paradoxus* reference strain CBS432 and extracted all the amino 5 acid changes. The unique amino acid change among the four parents are shown in red. We also 6 show *de novo* mutations identified in isogenic, two-parent and four-parent populations. We
1 predicted their functional impact by the online tool (mutfunc.com). The predicted scores are listed 2 in #Score and #IC columns. 3 The column description is as follows:

Given that the *TOR1, TOR2* standing and *de novo* variants often co-existed in the same population, we further investigated their potential interactions. We genotyped the local genetic background of *TOR1 de novo* mutant clones drawn from different endpoint populations and found their background genotypes can be different, i.e. the *TOR1 de novo* mutations had arisen on different clones (Table S4). Thus, *TOR1* mutations conferred strong adaptive gains across different clone backgrounds. This lack of a specific interplay between *TOR1 de novo* driver mutations and their genetic background is consistent with the observation that *TOR1 de novo* mutations also emerged and reached high frequency in all the isogenic populations. The interplay between *TOR2* mutations and their genetic background is particularly evident in a *TOR2* clone from population F12_2_RM_4. This clone carried the weak *TOR2^SA^* allele, whose frequency dropped from 0.29 (T0) to 0.04 (T8). However, the frequency was nevertheless enough for one *TOR2^SA^* clone to acquire a compensatory *TOR2* Ser1975Ile mutation in a heterozygote state. Consequently, the growth performance drastically increased and the *TOR2^SA^* allele frequency was driven to 0.46 at the end of selection (Figure S12B). Thus, the emergence of this *de novo TOR2* mutation in the *TOR2^SA^* allele compensated for the sensitivity of the *TOR2^SA^* allele by preventing RM binding. This indicated that standing and *de novo* variants of TOR probably act towards RM resistance via distinct mechanisms.

### Functional consequences of TOR natural variants

*TOR1* and *TOR2* are master regulators of growth, controlling yeast performance in many environments of relevance to industry, particularly in alcoholic beverage production. In this industrial context, we were particularly intrigued by the opposite RM resistance phenotype of the SA and WE *TOR1, TOR2* alleles, as they occur in two lineages independently domesticated for alcoholic beverage production (Fay & Benavides, 2005). We therefore further characterized the *TOR1* and *TOR2* alleles in these two genetic backgrounds to determine their respective impact on yeast performance in environments of industrial and medical interest. Potentially, such benefits could also explain their distinct evolutionary trajecTORIes.

First, given the role of TOR Complex 1 (TORC1) in regulating chronological life span (CLS) (Powers, 2006), we measured the impact of TOR variants on CLS in presence and absence of RM. In RM, the *TOR1^SA^* and *TOR2^WE^* alleles had antagonistic effects on birth and death rates, conferring faster growth and shorter CLS (Figure 5A-C). The wild type WE/SA carrying both copies of TOR had the shortest CLS and the shortest doubling time in RM, indicating *TOR1* and *TOR2* haplo-proficiency for CLS and haplo-insufficiency for growth in the hemizygous deletion strains. In the absence of RM, there was almost no difference in CLS between strains, indicating that the haplo-proficient effect of single copy *TOR1* and *TOR2* already saturated in rich synthetic medium.

Next, to understand the effects of *TOR1, TOR2* standing variants on TORC1 activity, we used a highly specific commercial antibody to measure phosphorylation of the ribosomal protein S6 (Rps6) under RM exposure. Rps6 phosphorylation is regulated by TORC1 and used as a specific *in vivo* assay for TORC1 activity (González et al., 2015). Rps6 phosphorylation increased in strains with the strong *TOR1^SA^* and *TOR2^WE^* alleles (Figure 5D). Thus, the SNPs distinguishing these alleles enhance TORC1 activity. This was quite surprising, first because a majority of SNPs in these alleles occur outside functional domains and second because not even mutations in the RM-binding domain affect TORC1 activity (González et al., 2015). This further underscored that the standing and *de novo* variants of *TOR1* and *TOR2* cause RM resistance by distinct mechanisms.

RM is an unlikely selection pressure on natural yeast alleles; however, real ecological constraints such as nitrogen limitation do affect cell growth in a TOR-dependent functions (Loewith & Hall, 2011) and is of central importance in wine fermentations. To further explore opposing *TOR1, TOR2* allele preferences between SA and WE backgrounds and illuminate the underlying mechanism, we measured their effect on doubling time in 18 relevant environments, including nitrogen-limitations and synthetic wine must (Figure 5E). As expected, the WE strain grew the fastest in synthetic wine must, consistent with its niche-specific domestication history. Overall, the removal of one *TOR* allele tended to result in growth defects in nitrogen-limited environments, and the removal of a WE allele was generally worse than the removal of a SA allele. For example, hybrids carrying *TOR1^WE^* grow faster than those carrying *TOR1^SA^* on methionine and threonine; and hybrids with *TOR2^WE^* grow faster than those with *TOR2^SA^* in tryptophan, threonine, serine, methionine, isoleucine, asparagine and adenine.

Finally, we investigated *TOR* gene essentiality in SA, WE, WA and NA genetic backgrounds by knocking out *TOR1* or *TOR2.* Previous studies in the laboratory strain S288C showed that *TOR1* was non-essential, whereas *TOR2* was essential (Liu et al., 2015; Winzeler et al., 1999). As expected, *TOR2* could not be deleted in WE, NA or WA. Surprisingly, however, we successfully deleted the *TOR2* gene in the SA haploid. The *tor2Δ* SA strain was able to grow on synthetic complete medium (SC), although with marked growth defects, but not on YPD (Figures 6A-B). Because the TORC1 activity of *tor2Δ* SA remained unaltered upon RM treatment (Figure 6C), the SA background is either able to make up for the *TOR2* loss by compensatory induction or by complex incorporation of *TOR1,* or do not use *TOR2* in TORC1 at all. We further dissected ∼900 spores from WE/SA *TOR2* reciprocal hemizygous deletions, as well as WE/SA wild type on both YPD and SC medium. On SC, the spore viability was 83.5% for the homozygote *TOR2* cross and 55.3% for the hemizygote cross. Thus, *TOR2* was essential in a fraction of the recombined WE/SA offspring. By tracking the deletion marker, we estimated that 18.5% of the *tor2Δ* spores carrying recombinants survived on SC (Table S7), although with large growth defects (Figure 6D). No *tor2Δ* spores were viable on YPD. Therefore, *TOR2* was conditionally essential, depending on both genetic background and growth condition. The conditional essential phenotype was usually regulated by complex genetic interactions, relying on multiple background-specific modifiers (Dowell et al., 2010). Tetrad segregation patterns suggested that there were at least two distinct loci contributing to *TOR2* dispensability (Table S8). Taken together, the divergent functions of the WE and SA alleles on *TOR1* and *TOR2* alleles impacted not only on RM resistance but also on chronological aging, TORC1 activity, nitrogen control of growth and essentiality. This may reflect the independent domestication hisTORIes of the two SA and WE lineages for specific purposes (Fay & Benavides, 2005).

**Figure 6.**
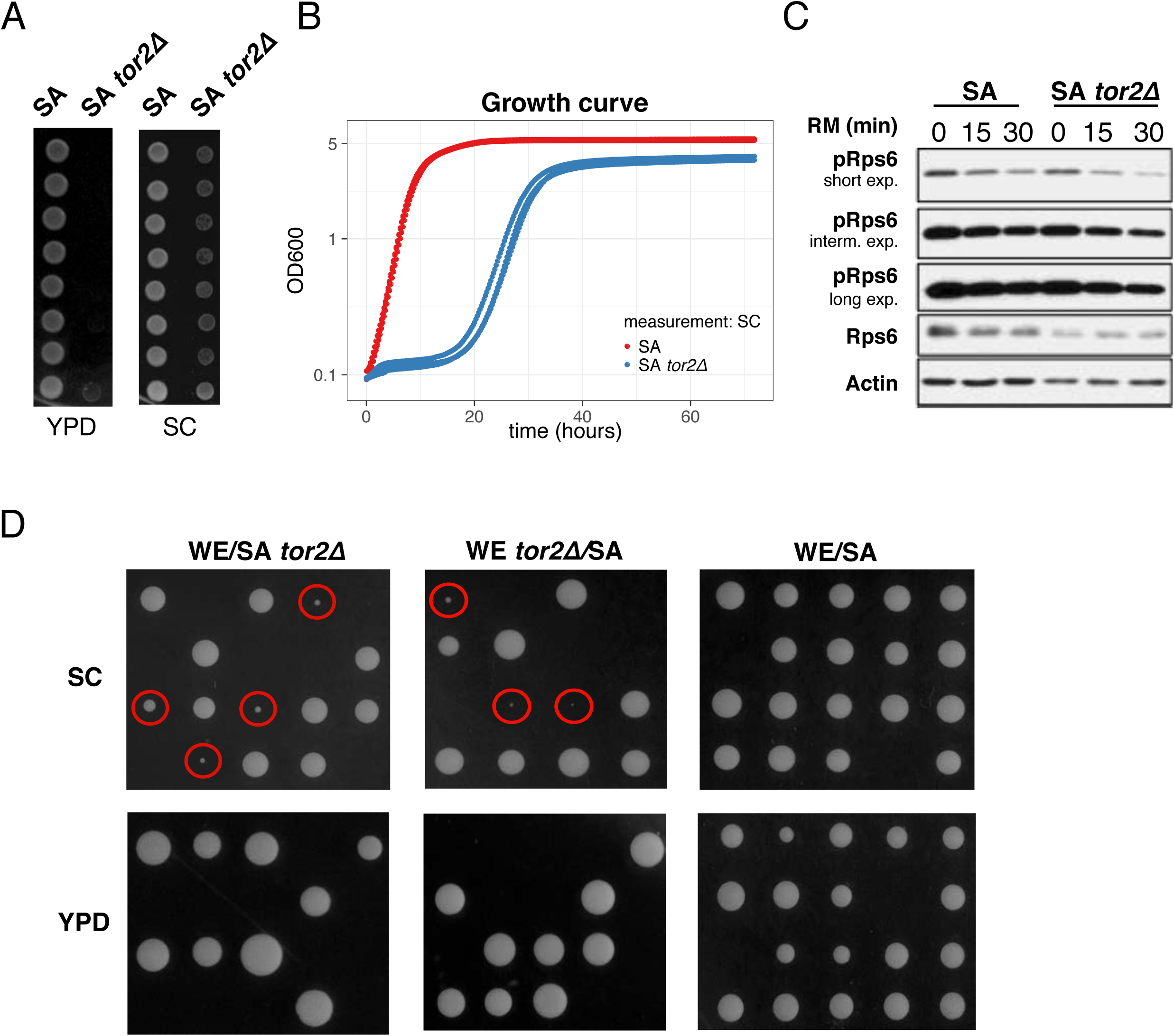
Functional characterization of the *TOR2* variants. (A) The SA *tor2Δ* cells are able to grow on synthetic complete (SC) medium although with visible growth defect, but not on YPD. (B) Growth curves of the SA *tor2Δ* and SA wild type strains in SC. (C) Immunoblot analysis showed Rps6 phosphorylation in SA wild type and *tor2Δ* strains shows that TORC1 activity is not altered by the *TOR2* deletion. All conditions are similar to the one reported in Figure 5D. (D) Representative plates acquired 4 days after tetrad dissection on SC and YPD for WE/SA wild type and its *TOR2* reciprocal hemizygotes. The red circles indicate viable *tor2Δ* strains.

## Discussion

We devised an experimental system with two (Parts et al., 2011) and four (Cubillos et al., 2013) parent intercrossed yeast populations to quantify how increasing levels of standing variation affect adaptation dynamics and to understand whether standing and *de novo* variants are selected in shared target genes. To maximize the genetic and phenotypic diversity, the four-parent populations were derived from intercrosses of four *S. cerevisiae* genetic backgrounds representative of independent evolutionary hisTORIes that were isolated from four different continents and ecological niches (Liti et al., 2009). The heterogeneous populations framework generated millions of individuals with unique haplotype combinations and has enabled high sensitivity and resolution QTL mapping (Burke et al., 2014; Cubillos et al., 2013; Illingworth, Parts, Schiffels, Liti, & Mustonen, 2012; Parts et al., 2011; Vázquez-García et al., 2017). Here the higher genetic heterogeneity translated into higher fitness variance which is a prerequisite for faster genetic adaptation (Jerison et al., 2017). The genetically diverse populations exploited on this variation to achieve larger and faster adaptive gains. We found that a doubling of the segregating genetic diversity (from 1/230 to 1/120 segregating sites/bp) increased RM adaptive gains by 51.5% and HU adaptive gains by 64.1% in the absence of *de novo* mutations. Undoubtedly, a continuum of genetic diversity and a large ensemble of environments are required for precise models of adaptation as a function of genetic variation. Nevertheless, these parameter estimates provide a starting point for placing the evolutionary theory of standing variation on a sound empirical basis. In terms of practical implications, this study also underscores the importance of minimizing the genetic variation of infections and tumors to maximize success rates when treating clonal evolutionary diseases.

Allele frequency dynamics in the four-parent populations revealed localized directional changes (QTLs) driving the early acceleration adaptation. The surprisingly high number of QTLs (13 in RM, 9 in HU) vastly exceeded the single QTL (*CTF8* in RM) mapped in the two-parent populations. The difference was partially a matter of new alleles being available in the four-parent populations: the largest effect QTLs (*RNR4, TOR1* and *TOR2)* were driven by WE and SA alleles that were not present in the two-parent populations. However, other four-parent QTLs corresponded to WA and NA alleles that were also present but not selected in the two-parent populations. Their lack of expressivity in the two-parent populations directly points to dependence on complex interactions conditioned by the higher genetic heterogeneity (Burke et al., 2010).

Towards the mid and later phase of selection, highly resistant clones emerged and arose to high frequency in both HU and RM. Nevertheless, the genetic make-up and origin of these clones differed dramatically between the two selection regimes. Standing variants appeared to drive HU adaptation all the way to the end, implying that beneficial *de novo* mutations are either too rare or too weak to compete against the bulk dynamics driven by the standing beneficial alleles (i.e. the nine QTLs). It could also be partially explained by negative or sign epistasis weakening the effects of beneficial *de novo* alleles (Khan, Dinh, Schneider, Lenski, & Cooper, 2011). An additional explanation is that the *RNR* driver mutations appear to be strongly background dependent. This is evident from the isogenic populations, with only one background (NA) that acquired *RNR4* mutations and evolved. In contrast, mid to late adaptation to RM was consistently driven by clones with *de novo* mutations in *TOR1, TOR2* and *FPR1* emerging and overtaking other competing bulk subpopulations. This was consistently true in all genetic contexts and at all levels of standing variation. These highly penetrant *de novo* mutations in members of the TOR pathway have long been known to prevent their interaction with RM (Heitman et al., 1991; Helliwell et al., 1994), which is manifested again by our experiments.

In the widest sense, we found strong examples of convergent selection on standing and *de novo* variants to confer RM resistance – *TOR1* and *TOR2.* This was not given *a priori.* First, strong loss-of-function *de novo* variants often play an outsized role under adaptation to a single constrained selective pressure (Hottes et al., 2013). However, such mutations are not likely to prevail in natural populations, because purifying selection acts to remove variants that impair gene functions (Bamshad & Wooding, 2003). Second, many standing variants in natural populations may not emerge *de novo* because the underlying mutation events are too rare. The convergence on selection on both standing and *de novo* variants of *TOR1* and *TOR2* is particularly intriguing. *De novo* and standing variants conferred RM resistance via distinct mechanisms: abolishing drug binding by *de novo* variants (Loewith & Hall, 2011) and altering the TORC1 activity by standing variants. Underscoring this mechanistic distinction, a driver mutation in the drug-binding domain completely rescued the low TORC1 activity of the weak *TOR2^SA^* allele. Moreover, the *TOR2 de novo* mutation is much rarer (only one single instance among all the populations of isogenic, two-parent and four-parent) despite its drug binding domain having a similar target size as *TOR1* and the *TOR1, TOR2* paralogs being thought to be redundant in terms of RM resistance (Loewith & Hall, 2011). The most parsimonious explanation for this drastic difference is that *TOR2* is under stronger selection constraints likely reflecting its unique, essential role in the TOR complex 2 (TORC2).

The standing WE and SA variants of *TOR1* and *TOR2* have opposite effects on RM resistance, reflecting lineage-specific functional divergence after the gene duplication in their shared ancestor. Although we cannot stringently reject a purely neutral explanation, the directly opposing effects of these alleles on growth and survival corresponds to an evolutionary trade-off between the two key determinants of fitness and makes it tempting to invoke selection to explain this divergence. Domestication to distinct human made niches (Sake and grape-wine), including different substrates of fermentation (Giudici & Zambonelli, 1992; Sasaki et al., 2014), may be the ultimate explanation for this divergence with drug resistance as a side-effect caused by TOR pleiotropy. More broadly, this is reminiscent of methicillin-resistant and penicillin-resistant strains emerging long before the introduction of these antibiotics in the clinic because of other irrelevant selections (Baker et al., 2014; D’Costa et al., 2011; Harkins et al., 2017).

Recent studies have implicated intratumoral heterogeneity as a significant driver of drug resistance, bearing big challenges to chemotherapy (Saunders et al., 2012). Both of the two key findings in this study: the acceleration of adaptation by higher standing variation; and the shared targets between standing and *de novo* variants have important implications on our understanding of drug resistance evolving and treatment development (McGranahan & Swanton, 2017). In particular, accurately measuring intratumoral heterogeneity and the clonal fitness distribution will become essential for more successful therapies in the near future.

## Materials and Methods

### Experimental evolution and genome sequencing

We previously performed two independent intercrosses to generate the F12 populations (four-parent populations – F12_1 and F12_2) that were derived from four diverged parents: DBVPG6044 (West Africa, “WA”), DBVPG6765 (Wine European, “WE”), Y12 (Sake, “SA”) and YPS128 (North America, “NA”). The strain information is listed in Table S9. Here experimental evolution was initiated from random subsamples of F12_1 and F12_2, with each subsample comprised of 10^7^ - 10^8^ cells. In parallel, experimental evolution was also initiated from clonally expanded, near isogenic parental populations of similar size. Cells were evenly spread on YPD agar plates (2% peptone, 1% yeast extract, 2% glucose, 2% agar) with hydroxyurea (10 mg/ml) or rapamycin (0.025 μg/ml), and incubated at 23°C. Every 2-3 days, all the cells were collected from each plate into 1 ml distilled water. Ten percent of the cell suspension was transferred to a freshly made plate while the rest were kept in 25% glycerol at -80°C. The selection experiment lasted for 32 days. The detailed timeline and population specifics are listed in Tables S1-S2. For each drug, there are four independently evolving replicates derived from F12_1 and F12_2 respectively, as well as two replicates for each of the four parents. Besides,there were two replicates derived from F12_1 and F12_2 respectively using drug-free YPD as control. Procedures were identical to those used for generating and evolving the previously published two-parent population (Vázquez-García et al., 2017). DNA was extracted from populations of T0, T1, T2, T4, T8 and the last transfer using “Yeast MasterPure” kit (Epicentre, USA). The samples were sequenced with Illumina TruSeq SBS v4 chemistry, using paired-end sequencing on Illumina HiSeq 2000/2500 at the Wellcome Trust Sanger Institute. Sequence data is deposited to NCBI SRA database with accession number for BioProject PRJEB4645.

### Sequence alignment, calling segregate genotypes and identification of *de novo* mutations

Short-read sequences were aligned to the *S. cerevisiae* S288C reference genome (Release R64-1-1). Sequence alignment was carried out with Stampy v1.0.23 (Lunter & Goodson, 2011) and local realignment using BWA v0.7.12 (Li & Durbin, 2009). We used SAMtools v1.2 (Li, 2011) to count the number of reads reporting parental alleles at the segregating sites (Cubillos et al., 2013). We performed *de novo* mutation calling for each sequenced sample using three different algorithms: GATK 2.1-5-gf3daab0 (DePristo et al., 2011), Platypus v0.7.9.1 (Rimmer et al., 2014) and SAMtools v1.2 (Li, 2011). We then filtered these calls by subtracting all variation called from the parental samples to remove standing variation, required each variant to be on a locus with more than ten reads and more than six reads reporting the variant allele, and to pass default filters of the algorithms. For Platypus we allowed allele bias flagged calls as the sequenced samples are pools and therefore can have a range of variant allele fractions. We then intersected the calls and required that at least two of the methods called it. For the confirmed driver mutations at the end time point, we further tracked their frequency across previous time points. Finally, we used Ensembl Variant Effect Predictor to annotate the mutations (McLaren et al., 2016).

### Estimating allele frequencies

We define the allele frequency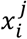at locus *i* of an allele *j* in the cross, e.g. we define 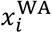 to refer to the frequency of the WA allele at locus *i* (and so on for *j* ∈ {WA, NA, WE, SA}). The allele frequency at locus *i* is normalized, such that 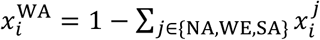. Given the number of reads 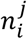 mapping to each allele and the total number of reads at each segregating locus, we estimated the allele frequency using the filterHD algorithm (Fischer, Vázquez-García, Illingworth, & Mustonen, 2014). filterHD fits a jump-diffusion process to the data where the diffusion component models the persistence of allele frequencies along the genome, reflecting linkage disequilibrium of nearby loci. Conversely, the jump component allows sudden changes in the allele frequency, which reflects the genotype state of large clones in populations that became clonal during the experiment.

### Estimating copy number variation

Sequencing depth was calculated by “samtools depth” and then used to calculate the median sequencing depth (*x*) for each chromosome. For the isogenic SA populations, we measured the z-score = (*x* - *μ*)/ *σ*, here *μ* and *σ* is the mean and standard deviation of sequencing depth of each population.

### Mapping quantitative trait loci (QTLs)

Given our allele frequency estimates, we used a 10-kb sliding window with a 2-kb step size to localize quantitative trait loci (QTLs). For each heterogeneous population, we compared the allele frequency change in a window *i* between time point *t* and T0 (e.g. 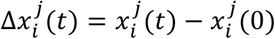, *j* = {WA, NA, WE, SA}). If there is selection on standing variation, the absolute frequency change of a parental allele in regions under selection is expected to be higher than in neutral regions and to increase gradually as selection proceeds. On this basis, for each earlier transfer, we calculated z-score of allele frequency changes compared with T0 in each population: 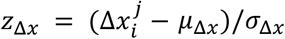. Here, *μ_Δ*x*_* and *σ_Δ*x*_* are the mean and standard deviation of 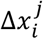 in all the four-parent populations evolved in the drug at a certain time point. The z-score square reflects the allele frequency deviation from T0. Given the fact that we observed dominant clones at later phase, we only used earlier phase to map QTLs: T0 to T4 for HU and T0 to T2 for RM. This cut-off is determined by the patterns of allele frequency distribution (Figure S8). Without dominant clone(s), the allele frequency distribution of all the four parental lineages follows a normal distribution with mean of ∼0.25. If dominant clone(s) appear and greatly deplete the genetic heterogeneity of the population, the distribution pattern would change dramatically, such as the ones shown at later time points with two or more peaks of frequencies largely deviated from 0.25. We searched for regions with z-score square higher than 99% or 95% quantile for each earlier time point. If the regions were able to pass the cut-off at T1, T2 for RM and at T1, T2, T4 for HU, and not pass the same cut-off in control (drug-free condition), they are assumed to be QTLs (Figures 4A and S10, Table 1). We excluded regions located near chromosome ends, which could be false positives due to repetitive sequences. The discrepancy of QTL numbers between the two-parent and the four-parent populations cannot be attributed to the different approaches to perform QTL mapping because when applying the same approach described here to the two-parent data, only few weak QTLs were mapped (Figure S13) including *CTF8.*

QTLs could be either maintained until later time points or be hijacked by the spread of clones with beneficial mutations. We define whether a QTL is maintained by counting the replicates in which the strong allele keeps increasing or the weak allele keeps decreasing until T4, T8 and the end. If the number of such replicates is more than six (of a total of eight), we defined the QTL as maintained until the later time points (Figure S9, Tables 1 and S5).

### Growth phenotyping

#### Quantitative measurement

We randomly selected thousands of isolates from the initial and final populations (Table S2), bulk population from the isogenic, two-parent and four-parent populations at each serial transfer of the experimental evolution (Table S1) and strains with gene deletion (Table S9) for phenotyping. Using a high-resolution large-scale scanning platform, Scan-o-matic, we monitored growth in a 1536-colony design on solid agar plate (Zackrisson et al., 2016). High-quality desktop scanners monitored the colonies growth on synthetic complete medium (0.14% YNB, 0.5% ammonium sulphate, 0.077% Complete Supplement Mixture (CSM, ForMedium), 2% (w/v) glucose and pH buffered to 5.8 with 1% (w/v) succinic acid) with drugs (10 mg/ml hydroxyurea, 0.025 ug/ml rapamycin), and without drug as control. The medium of nitrogen-limited environments to test the TOR variants used a single nitrogen source present at 30 mg nitrogen/l (Ibstedt et al., 2015). Experiments were run for 3 days and scans were continuously performed every 20 minutes. After filtering steps for quality check, doubling time was extracted for downstream analysis in R (R version 3.4.1). Technical replicates (n) are substantial: *n* ≥ 8 for each sample in drug condition; *n* ≥ 2 in drug-free condition; *n* ≥ 96 for the samples phenotyped in nitrogen-limited conditions.

We also used the Tecan Infinite 200 PRO plate reader to measure growth curves in small scale. We pre-cultured the cells overnight and diluted the saturated culture 100 times into fresh medium. We measured OD_600_ every 15 minutes for at least 3 days in drugs and control. The raw OD_600_ values were corrected and then used to generate growth curves. Doubling time and yield were extracted using the online tool “PRECOG” (Fernandez-Ricaud, Kourtchenko, Zackrisson, Warringer, & Blomberg, 2016).

#### Qualitative measurement

We did serial dilution and spotting of the cells to visualize the adaptation on population level visually (Figure S1) as well as the growth phenotypes of gene deletions (Figure S5). Cells were pre-cultured in YPD overnight to saturation. Then 5 μl of the culture was taken to do spotting assay in the condition of interest. There were a total of six 1:10 dilutions from left to right on the plate.

We also did spotting assay of 48 isolates drawn from the SA population evolved in RM (SA_RM_2_T15) in heat condition (40°C). We pre-cultured cells in YPD overnight. Then 5 μl cells of 1,000-fold dilution from saturation were taken to put on YPD and incubated at 40°C. The plates were scanned after two days.

### Chronological Life Span (CLS) measurement

Strains used for CLS measurement were thawed from -80°C and grown on YPD plate. Single colonies were picked and pre-cultured in 1 ml synthetic complete (SC) medium (0.675% YNB, 0.0875% complete powder, 2% glucose) overnight until saturation. Then the cells were mixed well and 5 μl overnight culture was transferred to 200 μl SC and 200 μl SC + rapamycin (0.025 μg/ml) in 96-well plate, which was sealed with aluminum foil and kept in an incubator at 30 °C. Each strain has four replicates. After 3 days, red fluorescent dye propidium iodide (PI) was used to stain the dead cells and green fluorescent dye YO-PRO was used to stain apoptotic cells. Double staining dyes were diluted in PBS at a final concentration of 3 μM for PI and 200nM for YO-PRO. Cells were well suspended by pipetting and 5 μl culture was transferred to 100 μl PBS with PI and YO-PRO, stained at 30°C for 10 minutes. Flow cytometry analysis was performed on the BD FACSCalibur system. Excitation was performed using a laser at 488 nm and emission was detected in FL1 and FL3 using the standard filter configuration. The first measurement was termed as Day0. After that, every 3-4 days, we used the same protocol to stain cells and measure viability.

### Quantitative PCR (qPCR) to confirm the chrIX copy number variation

In order to validate the chrIX copy number changes of the SA clones evolved from RM evolved population, we performed qPCR with StepOnePlus™ Real-Time PCR System. Primers were designed on both sides of the chrIX centromere to validate chrIX copy number changes. Another pair of primers was designed within an essential gene located on chrI (Table S10) as control. We made a standard curve (*R*^2^ = 0.99) for each pair of primers and melting curves of each qPCR product to make sure of amplifications specificity. DNA template was prepared using “Yeast MasterPure” kit (Epicentre, USA). Each qPCR reaction has three replicates using the FastStart Universal SYBR Green Master (Rox). We also used the SA wild type diploid as control. We used ΔΔCt method (Schmittgen & Livak, 2008) to analyze data to determine whether there are chrIX copy number changes.

### Cross grid experiment

We isolated diploid SA clones from a RM-evolved population at T15. We validated the copy number of chrIX (three or four copies), induced sporulation (in 2% KAc) and dissect spores. We genotyped the spores of the mating type, *TOR1* mutation or wild type and chrIX copy number (one or two copies). With these genotypes, we crossed spores to create an array of diploids where all possible genotypes were combined (Figure 2D).

### Reciprocal hemizygosity

Gene deletion was performed using LiAc/SS carrier DNA/PEG method (Gietz & Schiestl, 2007). Reciprocal hemizygosity analysis (Warringer et al., 2017) was performed in the hybrids derived from two of the four parents. We tried several times to construct *RNR4* reciprocal hemizygotes in the hybrids with WE allele (WE/NA, WE/WA and WE/SA) to confirm the function of strong allele (WE). Finally, we obtained complete reciprocal hemizygotes in WE/NA but incomplete in WE/WA and WE/SA (only WE allele deleted, but not WA or SA allele deleted). We successfully constructed complete reciprocal hemizygotes for *TOR1* and *TOR2* in the WE/SA hybrid (Table S9).

### Measurement of TOR activity by Rps6 phosphorylation

Exponentially growing cells (OD_600_ 0.6-0.8) in SC medium were treated with rapamycin (LC laboraTORIes) to a final concentration of 200 ng/ml. Cultures (10 ml) were centrifuged at 1800g for 2 min at 4°C. The cell pellet was washed once with 500 μl cold water and stored at -80°C. Protein extraction, SDS-PAGE separation and immunoblot analyses were performed as previously described (González et al., 2015). The antibodies used in this study are: phospho-Ser235/Ser236-S6 (#2211, Cell Signaling Technology), RPS6 (#ab40820, Abcam), actin (#MAB1501, Millipore).

### Tetrad analysis

Cells were sporulated in 2% KAc at 23°C. When at least 30% tetrads were observed under the microscope, we treated the cells in zymolase (5 mg/ml) at 30°C for 30 minutes. Then spores were dissected manually by the Singer SporePlay+ instrument. To validate the conditional essentiality of *TOR2*, both YPD medium (2% peptone, 1% yeast extract, 2% glucose, 2% agar) and SC medium (0.675% YNB, 0.0875% complete powder, 2% glucose, 2% agar) medium were used for tetrad analysis. Plates were photographed after 4 days and replica plated on SC + Nourseothricin to follow the segregation patterns of knockout alleles. The absence of *TOR2* in small spores is also confirmed by PCR. The statistical approach to identify modifiers for the conditional essentiality of *TOR2* is performed based on the method described by Dowell *et. al.* (Dowell et al., 2010).

### Sequence analysis of >1,000 yeast strains and function predictions

We reconstructed diploid pseudo-genome sequences of the 1,011 *S. cerevisiae* natural isolates by substituting the reference *S. cerevisiae* genome with the SNP calling results of the 1002 *S. cerevisiae* Genomes Project (Peter and De Chiara et al. under review). In the occurrence of heterozygous SNPs, we randomly distributed the two alleles into the two haploid pseudo-genomes. The first haploid pseudo-genome of each isolates was used for our downstream analysis. We extracted the CDS regions of the *RNR4, FPR1, TOR1* and *TOR2* genes from these haploid pseudo-genome sequences based on the reference coding-region coordinates and performed sliding window analysis (window size = 60 bp, step size = 0) for the coding region of each gene to calculate the pairwise sequence diversity (π) with Jukes-Cantor correction. Likewise, we also calculated the ratio of non-synonymous and synonymous substitutions (dN/dS) for each window using the “yn00” program of the PAML package (version 4.8a) (Yang, 2007). For dN/dS calculation, we used the corresponding sequences of the *S. paradoxus* strain CBS432 as outgroup. The coding sequences of those four genes in CBS432 were retrieved from our previous study (Yue et al., 2017) and were aligned with their counterparts of the *S. cerevisiae* strain genomes in codon spaces using MEGA7 (Kumar, Stecher, & Tamura, 2016) with indels trimmed off. All the amino acids substitutions from the 1,011 strains were submitted online to predict the functional consequences (mutfunc.com). The Tor1, Tor2 and Rnr4 protein sequences were analyzed by SIFT (Sim et al., 2012) and the alignments were used to calculate sequence conservation (Capra & Singh, 2007).

### Statistical analysis

The Mann–Whitney U-test was performed in R using the *wilcox.test ()* function, with two-sided alternative hypothesis. Unless otherwise stated, the doubling time mentioned in the text corresponds to the mean value of indicated samples.

## Acknowledgments

We thank Johan Hallin for critical reading of the manuscript. This research is supported by ATIP-Avenir (CNRS/INSERM), Fondation ARC (SFI20111203947), FP7-PEOPLE-2012-CIG (322035), the French National Research Agency (ANR-13-BSV6-0006-01 and ANR-16-CE12-0019), Cancéropôle PACA (AAP emergence) and DuPont Young Professor Award to G.L., by the Wellcome Trust to I.V.-G. (WT097678) and to V.M. (WT098051), by the Swedish Research Council (325-2014-6547 and 621-2014-4605) to J.W. J.L. is supported by Fondation ARC pour la Recherche sur le Cancer (PDF20140601375). J.-X.Y. is supported by Fondation ARC pour la Recherche sur le Cancer (PDF20150602803). B.B. was supported by La Ligue Contre le Cancer (GB-MA-CD-11287). We also acknowledge the IRCAN Flow Cytometry Facility CytoMed (supported by Conseil Général 06, FEDER, Ministère de l’Enseignement Supérieur, Région Provence Alpes-Côte d’Azur and INSERM) and the IRCAN Genomics Core Facility.

## Supplementary Information

**Figure S1.**
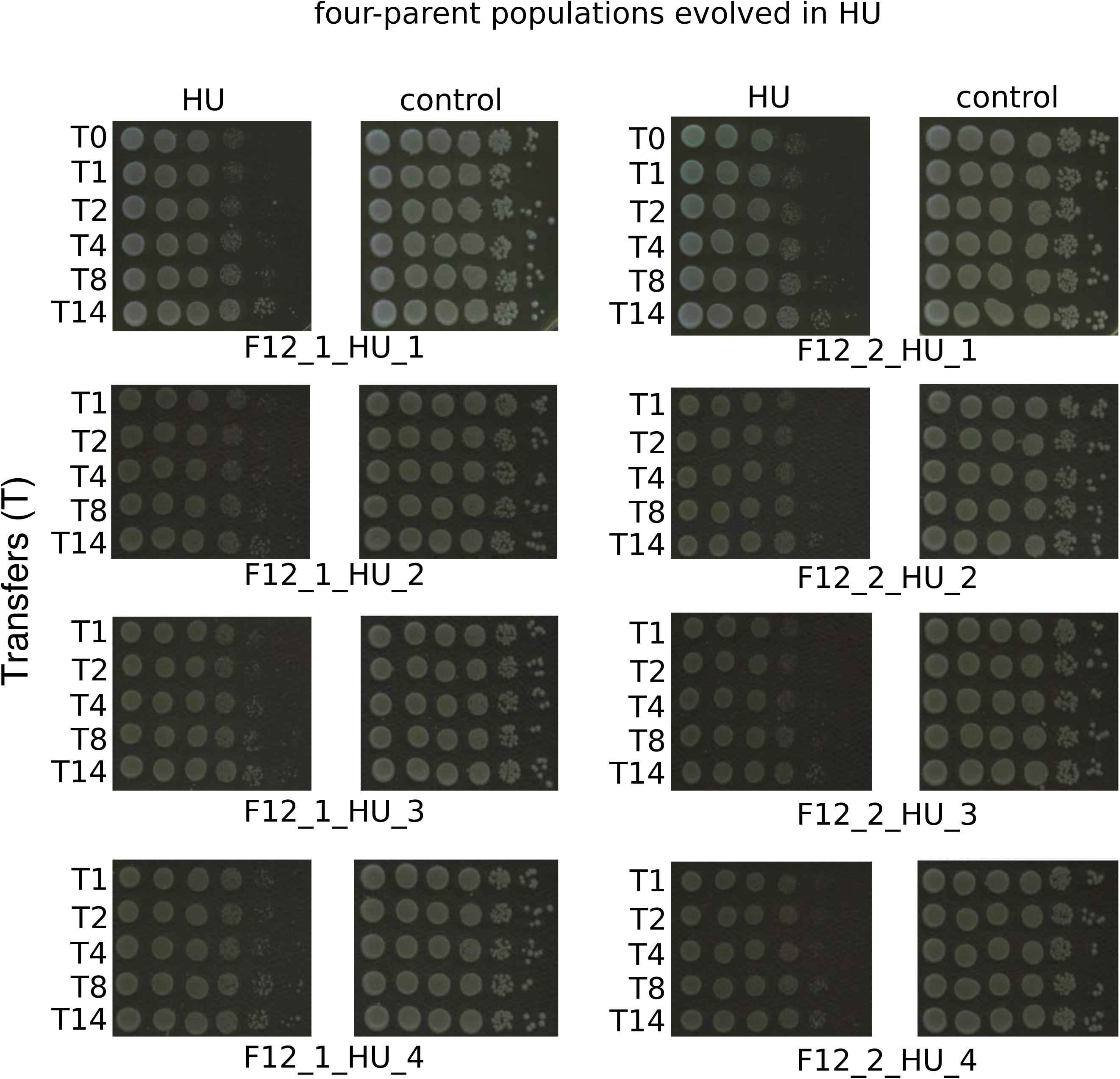
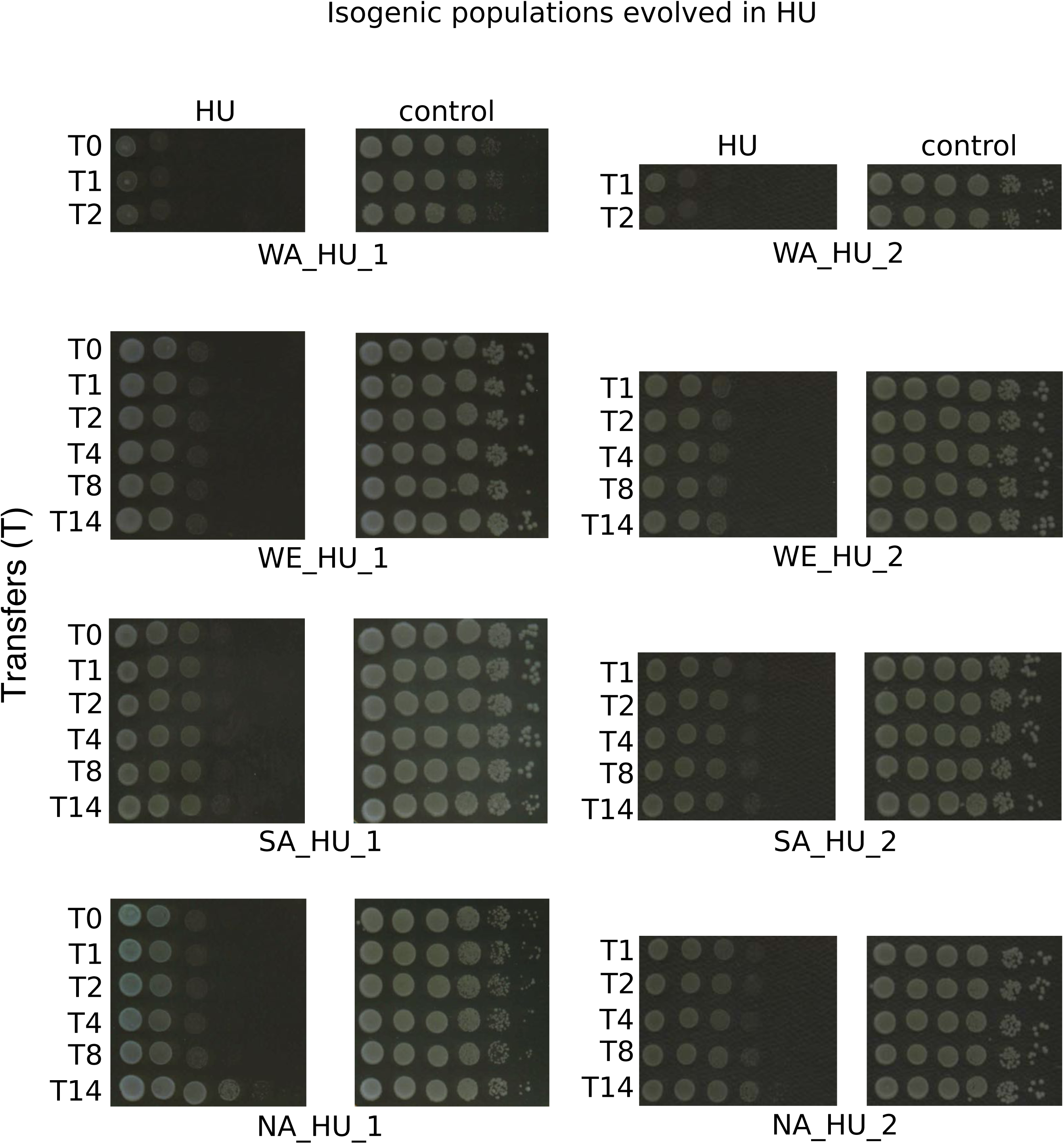
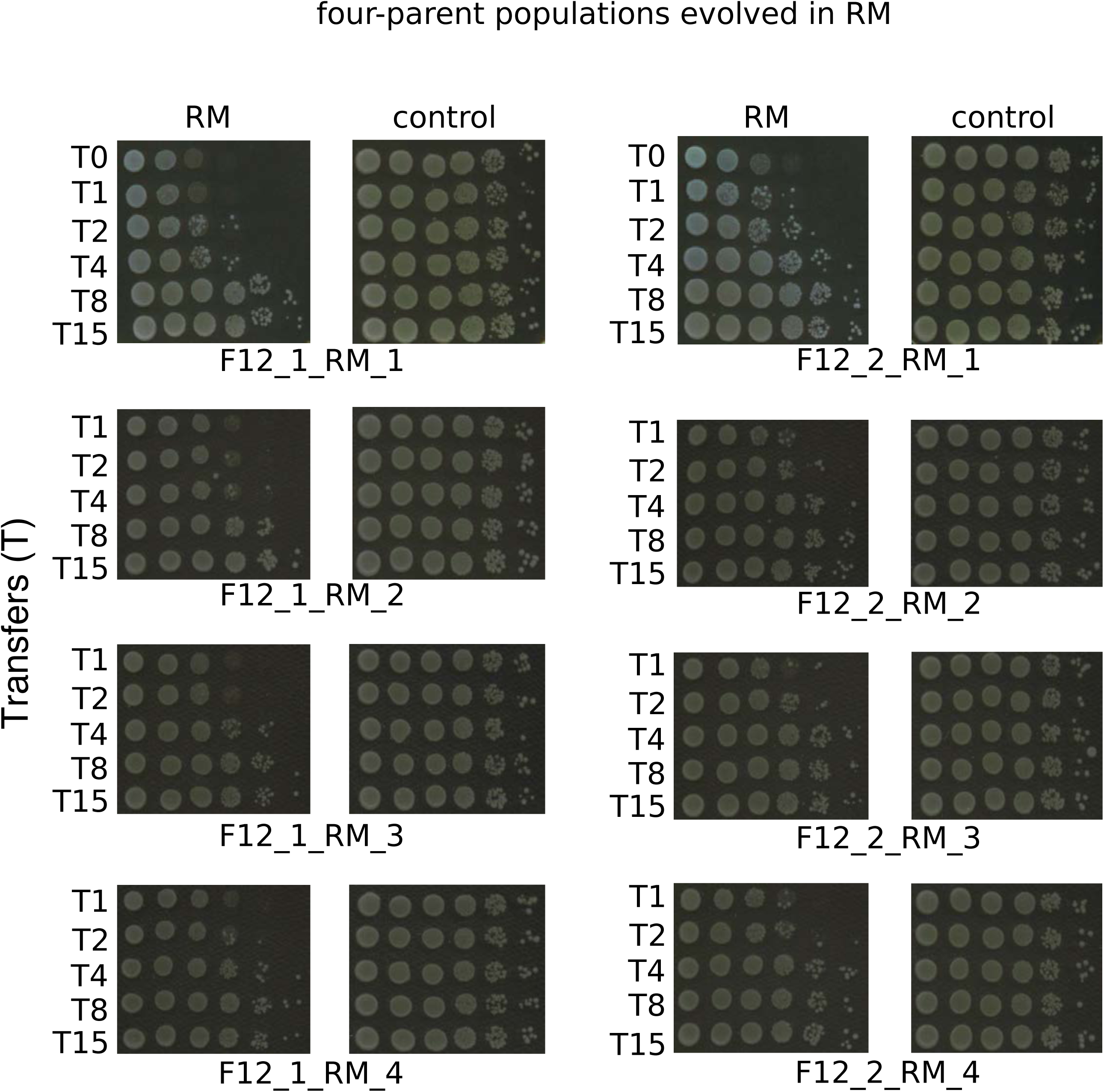
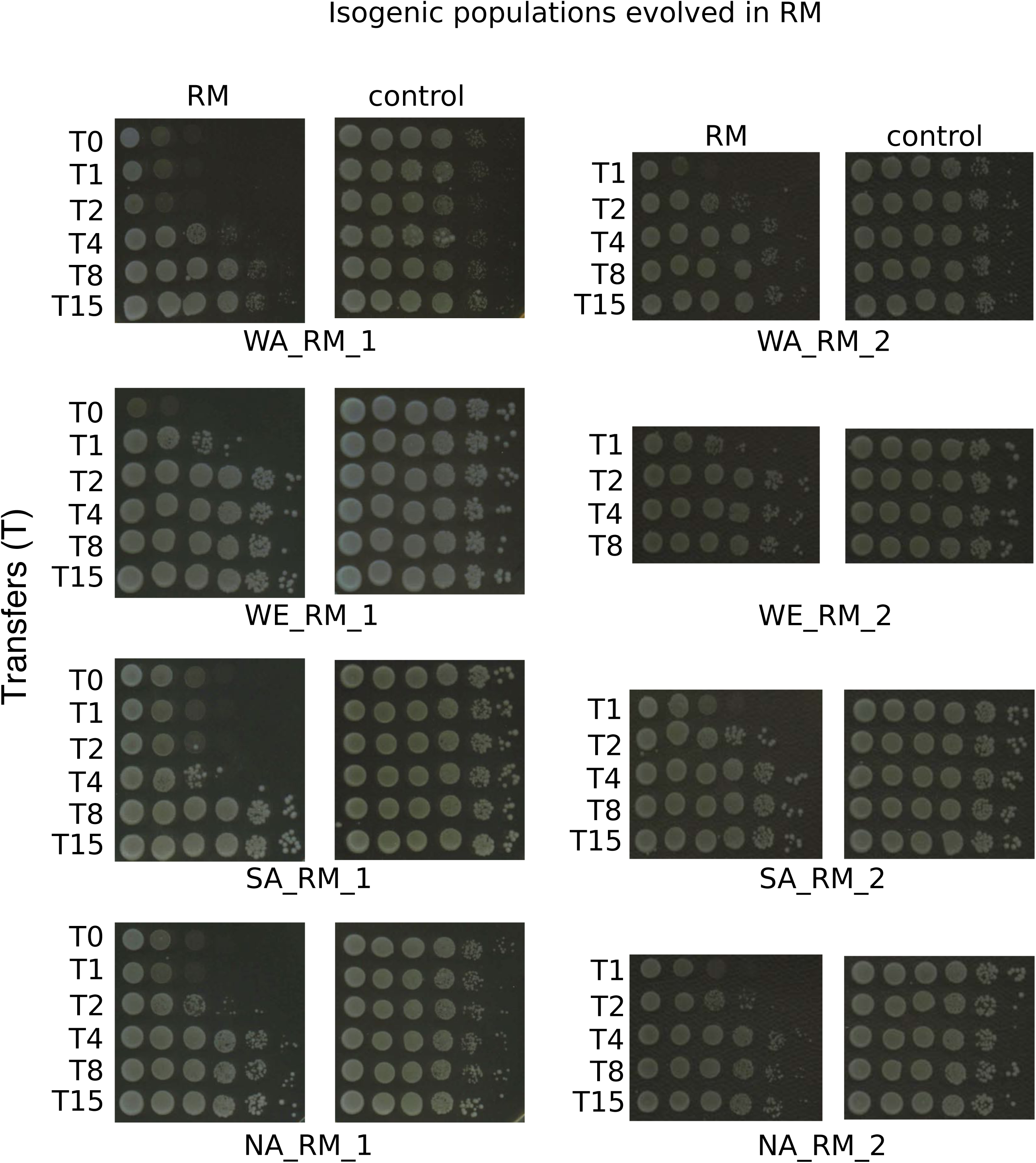
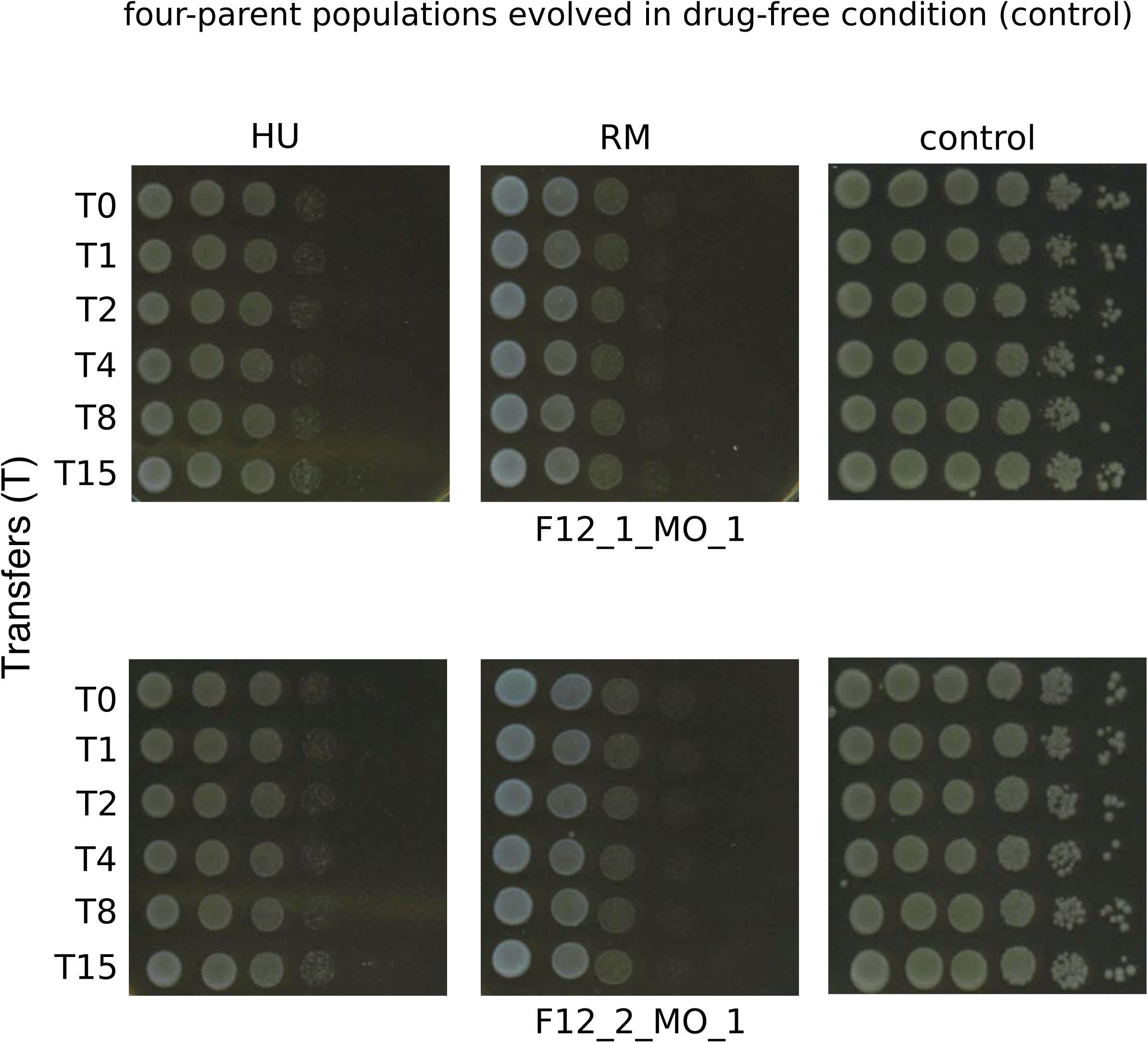
Spotting assay of all the four-parent and isogenic populations in this study. Populations were sampled at T0, T1, T2, T4, T8 and T14 (HU) or T15 (RM and control). Each spot represents 10-fold serial dilution of the cells from left to right.

**Figure S2.**
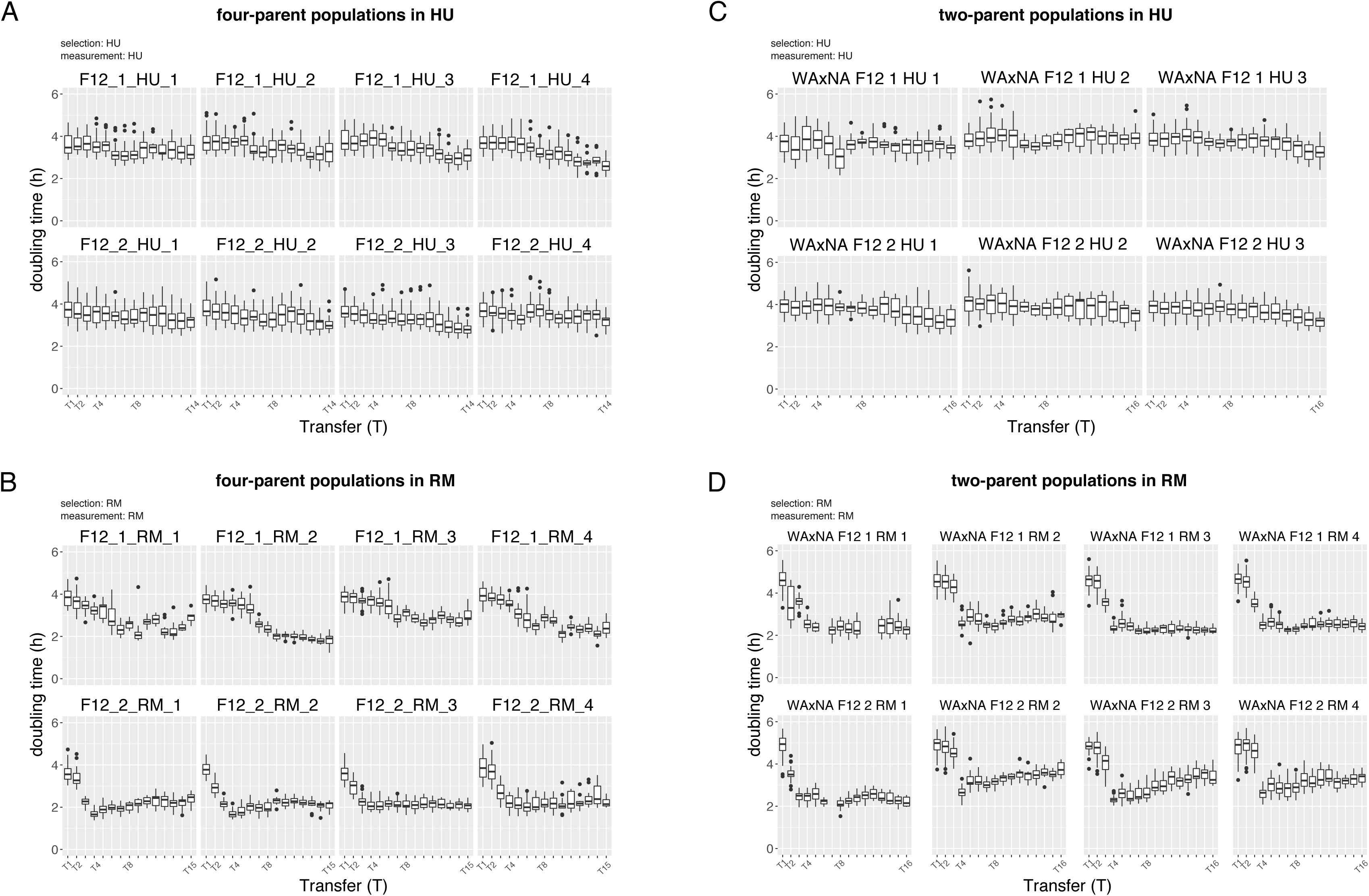
Doubling time of each randomly sampled heterogeneous bulk populations. (A)the four-parent populations in HU, (B) the four-parent populations in RM, (C) the two-parent populations in HU and (D) the two-parent populations in RM after each expansion cycle. The timeline for the experiment evolution is listed in Table S1. Boxplot shows the doubling time of all the technical replicates. Boxplot: Center lines, median; boxes, interquartile range (IQR); whiskers, 1.5×IQR. Data points beyond the whiskers are outliers.

**Figure S3.**
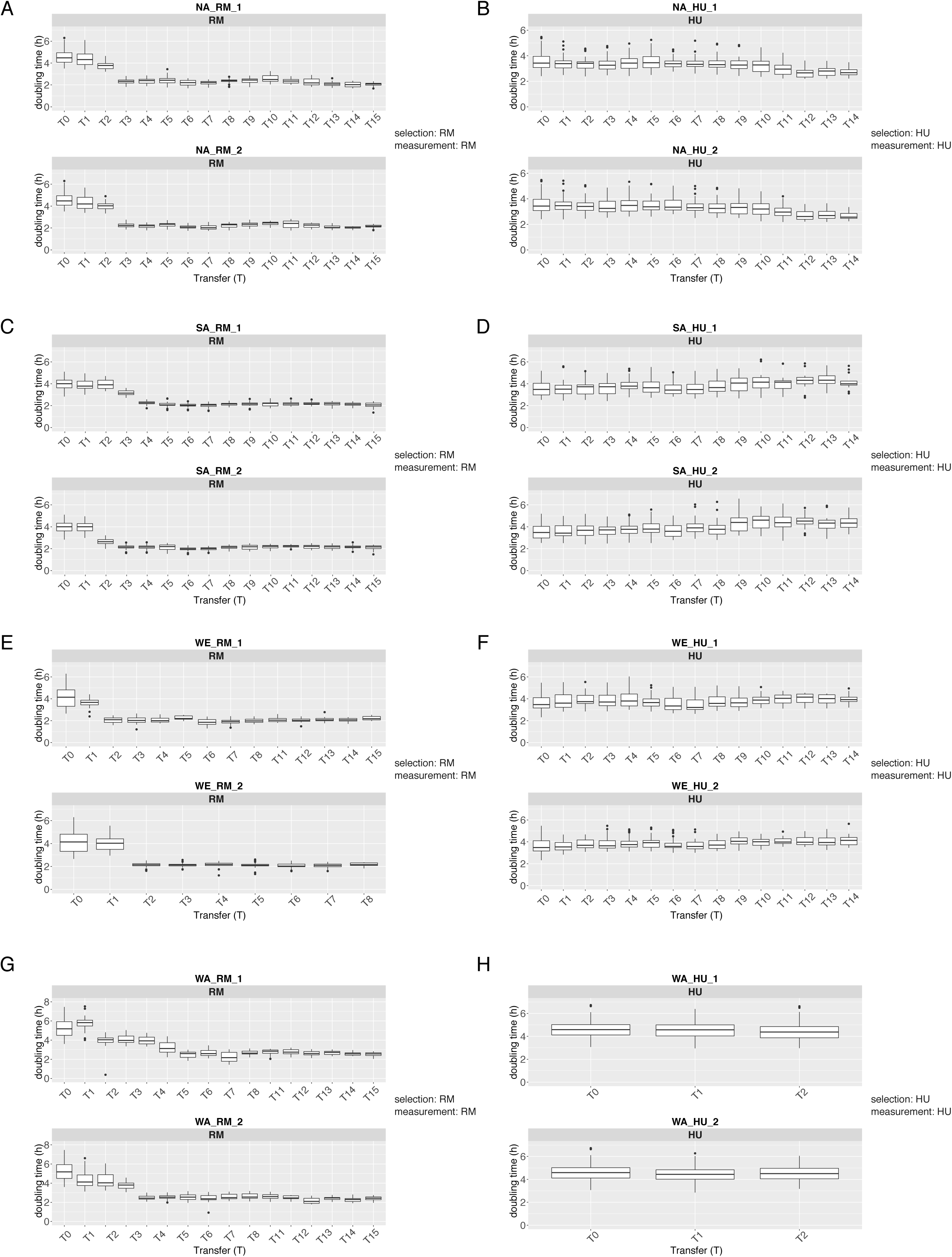
Doubling time of each randomly sampled isogenic bulk populations. Doubling time of isogenic populations evolved in RM (A, C, E, G) and HU (B, D, F, H). The top and bottom panels show two replicates of each parent. The timeline for the experiment evolution is listed in Table S1. WA in HU died out after T2. The second replicate of WE in RM was contaminated after T8. Boxplot shows the doubling time of all the technical replicates. Boxplot: Center lines, median; boxes, interquartile range (IQR); whiskers, 1.5×IQR. Data points beyond the whiskers are outliers.

**Figure S4.**
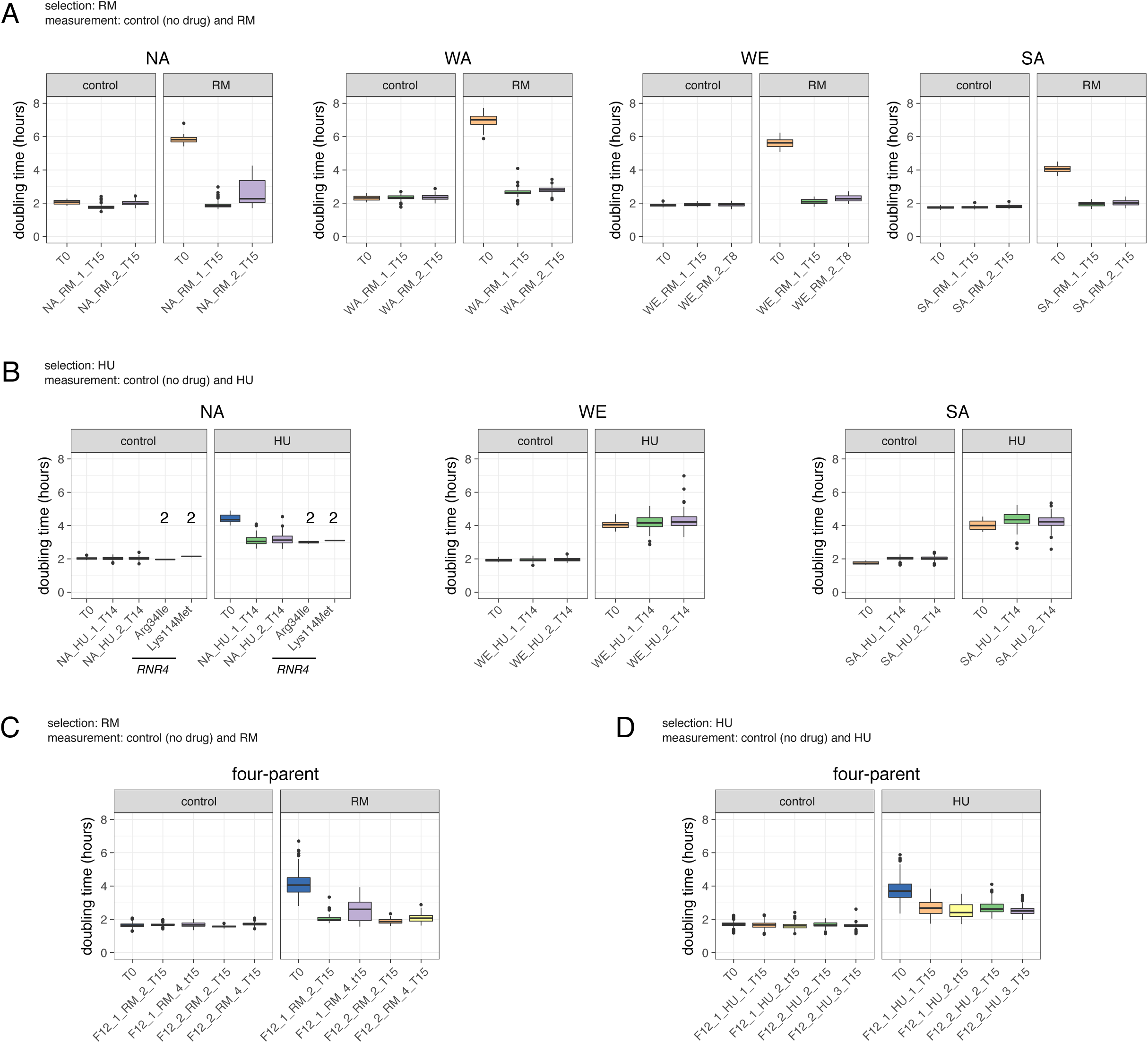
Doubling time of individuals drawn from the initial and final populations. Doubling time of individuals from (A) isogenic populations evolved in RM, (B) isogenic populations evolved in HU, (C) four-parent populations evolved in RM, (D) four-parent populations evolved in HU condition. There are 48 isolates from each ancestral and 96 isolates from each final replicate population. WA in HU is not included due to the extinction after T2. The doubling time of *RNR4* mutants from the NA populations is shown in (B, the NA panel). The number above the boxplot indicates the number of genotyped individuals with confirmed driver mutations by Sanger sequencing. Boxplot: Center lines, median; boxes, interquartile range (IQR); whiskers, 1.5×IQR. Data points beyond the whiskers are outliers.

**Figure S5.**
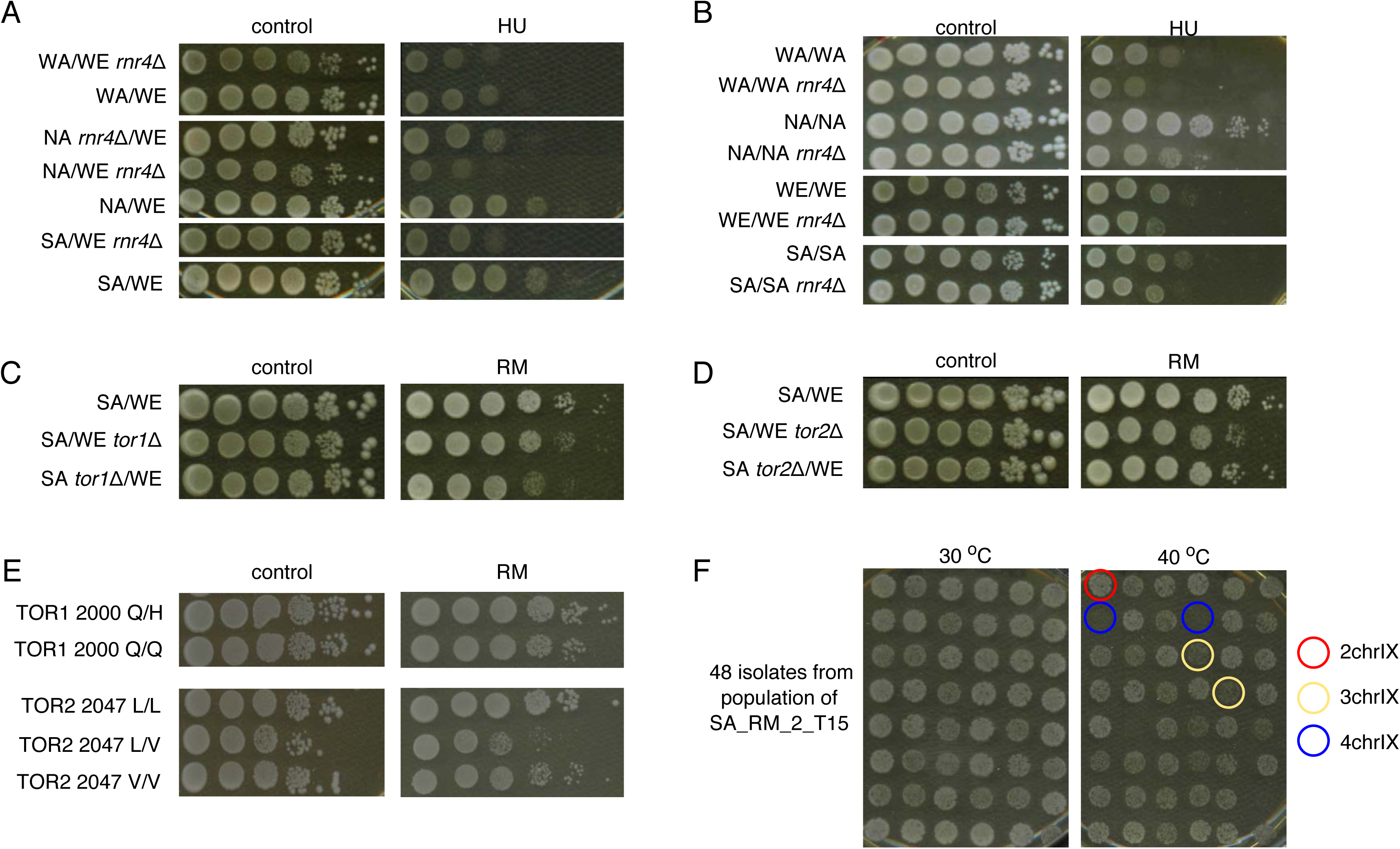
Spotting assay of the genetic constructs and strains from the 1002 Yeast Genomes project. (A) Phenotype of *RNR4* reciprocal hemizygosity constructs in HU and control. (B) Phenotype of *RNR4* hemizygous deletion of the four parental diploid strains in HU and control. Phenotype of (C) *TOR1* and (D) *TOR2* reciprocal hemizygosity constructs in RM and control. (E) Phenotypes of wild type strains from the 1002 Yeast Genomes project. The diploid strains with heterozygous Q/H amino acids at position 2000 in *TOR1* and homozygous L/L amino acids at position 2047 in *TOR2* show RM resistance. (F) Spotting assays of 48 isolates from SA endpoint population evolved in RM (SA_RM_2_T15) in heat condition (40°C). The isolates with red, yellow and blue circles were confirmed to have two, three and four copies of chromosome IX respectively by real-time PCR. Extra copies of chromosome IX lead to heat sensitivity.

**Figure S6.**
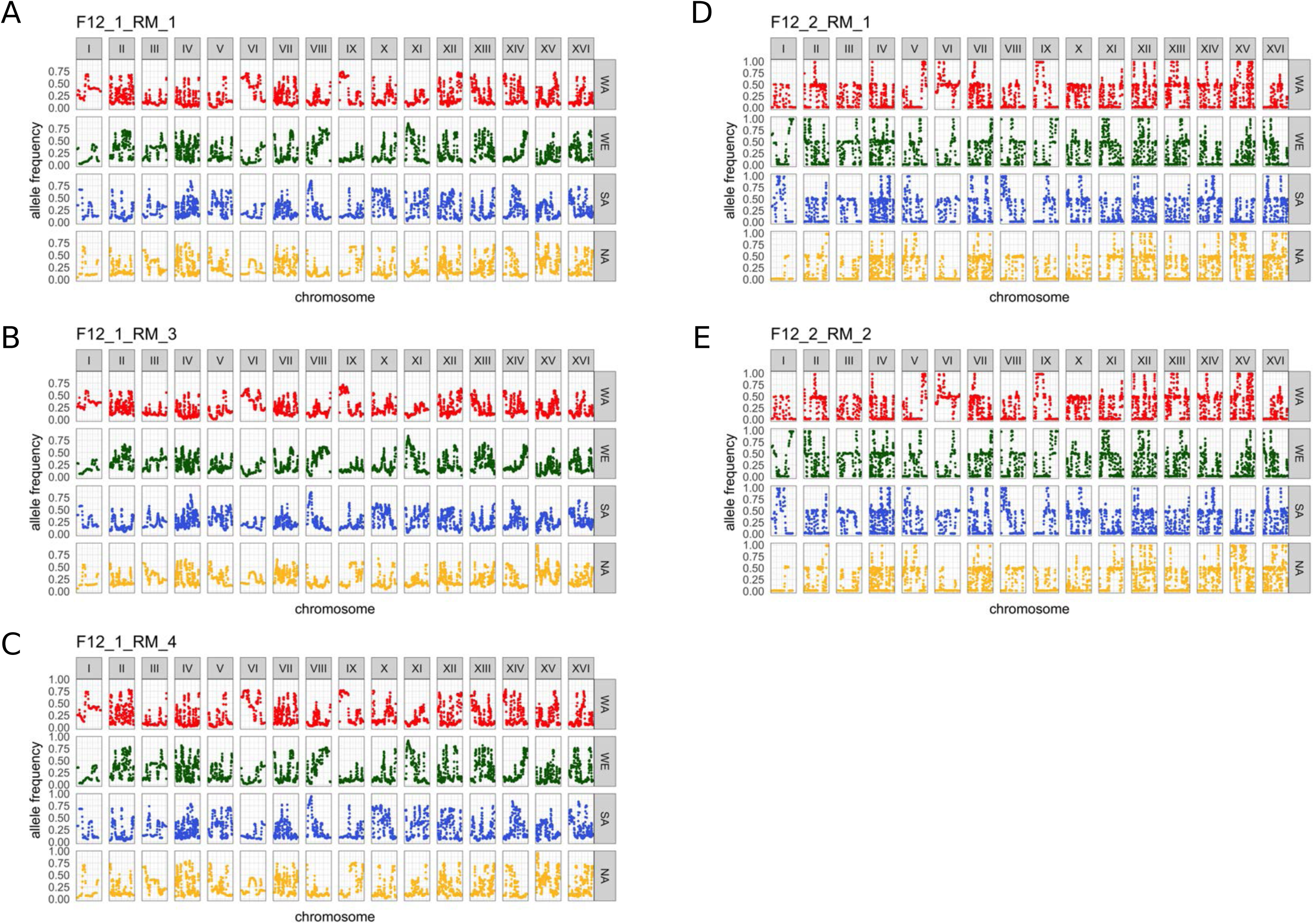
Genome-wide allele frequency of the endpoint populations. The endpoint populations (T15) evolved in RM shows similar pattern in replicates within the same intercross replica (F12_1 or F12_2) but different between them. We observed similar allele frequency pattern in populations that had the *FPR1* Thr82Pro mutation (A-C, F12_1 replicates); and that had the *TOR1* Trp2038Ser mutation (D-E, F12_2 replicates).

**Figure S7.**
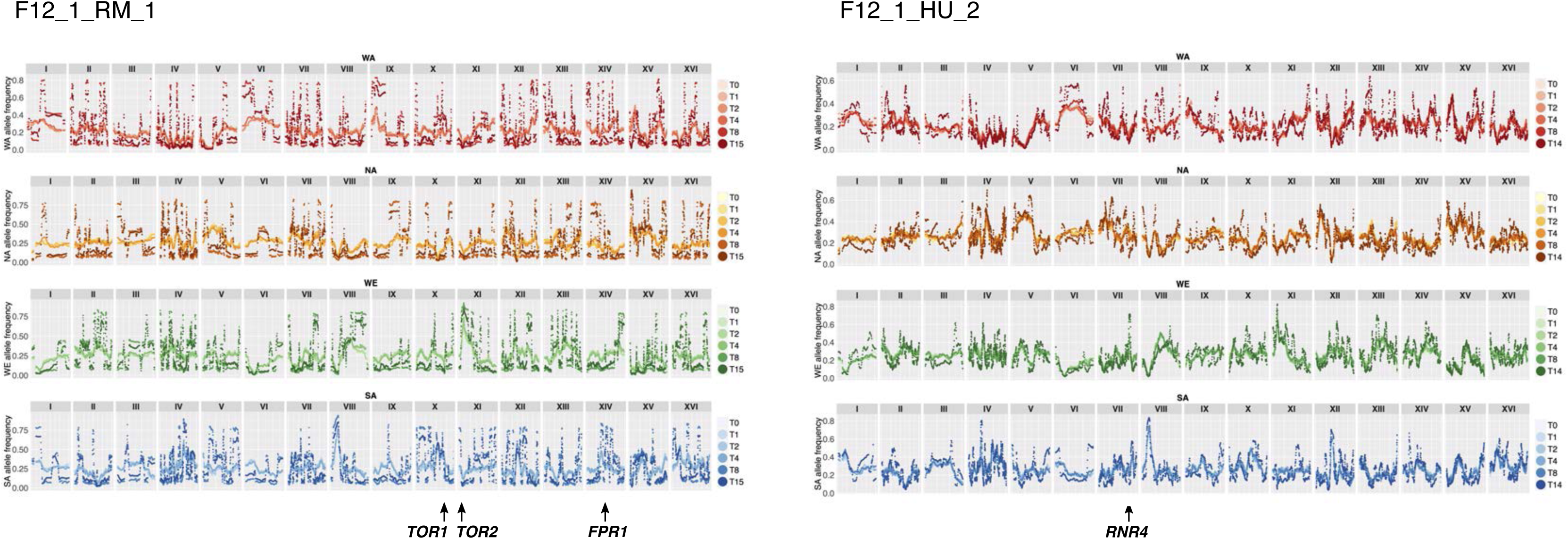
Genome-wide allele frequency changes across multiple time points. Two representative populations evolved in RM (left) and HU (right) are shown. The four panels show allele frequency changes of the four parental lineages (WA, NA, WE and SA from top to bottom). Early and late time points are indicated by colors from light to dark. At earlier phase, local allele frequency changes are the signal of selection on standing variation underlying responsible QTLs. At later phase, with the emergence of highly resistant clones, broad jumps in allele frequency are observed across the genome.

**Figure S8.**
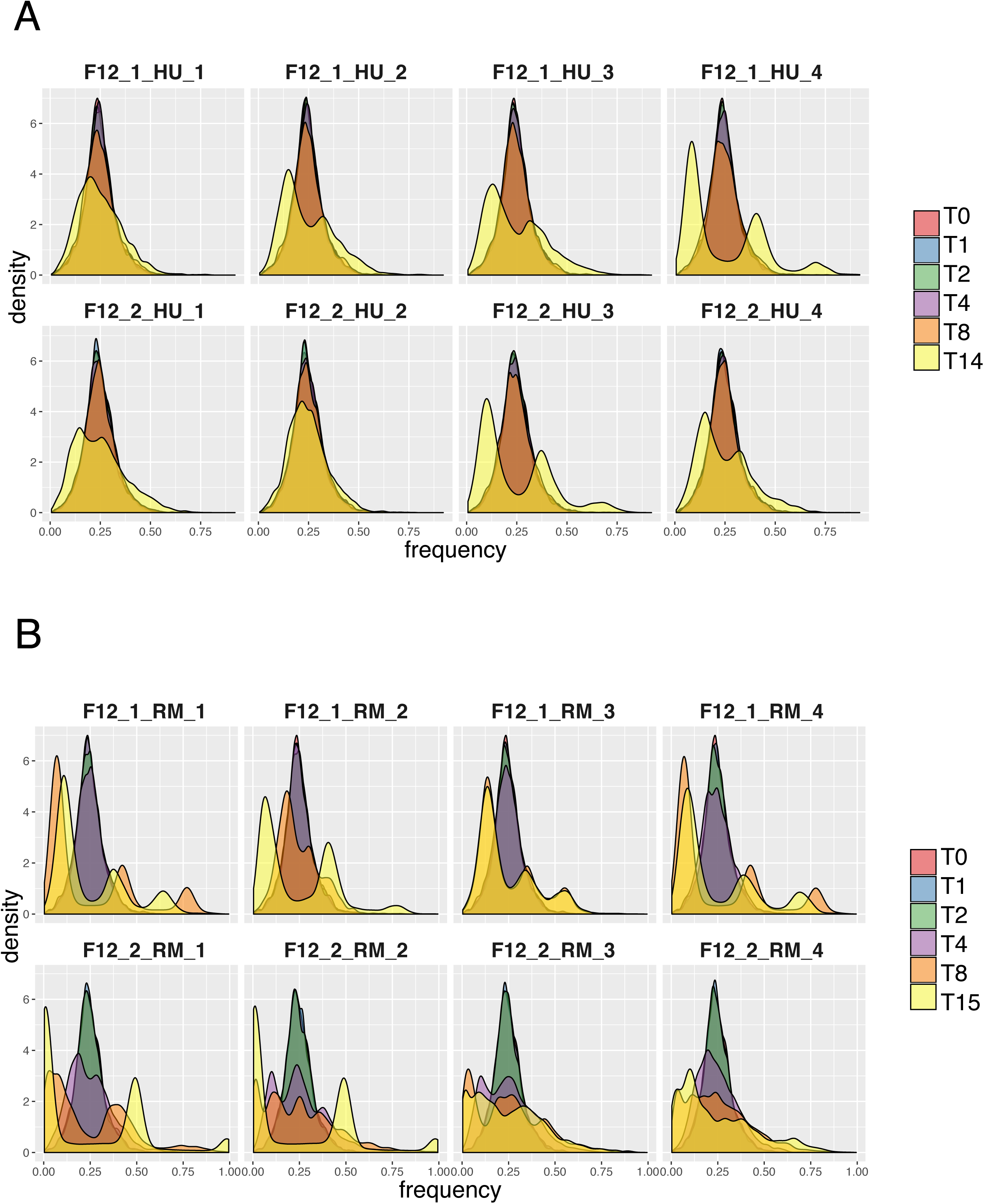
Four-parent population allele frequency density plots. Allele frequency distribution in HU (A) and RM (B) in all the replicates. Initially, the distribution of allele frequency derived from the four parents is close to a normal distribution centered on 0.25. The distribution pattern changes in late phase due to the emergence of resistant clones. To be conservative, we used T0 to T2 in RM and T0 to T4 in HU for the identification of QTLs, when selection mostly acted on standing variation.

**Figure S9.**
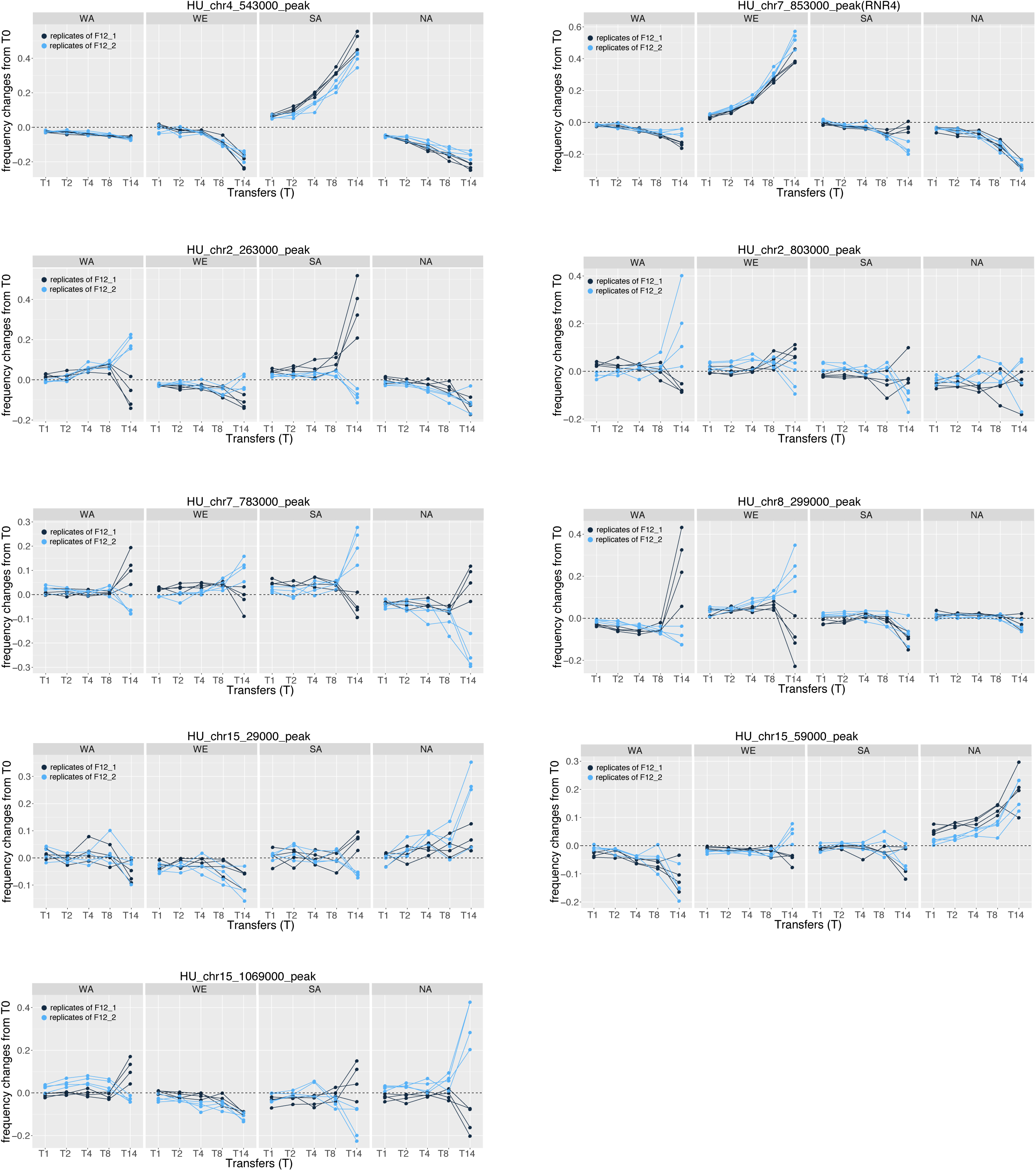
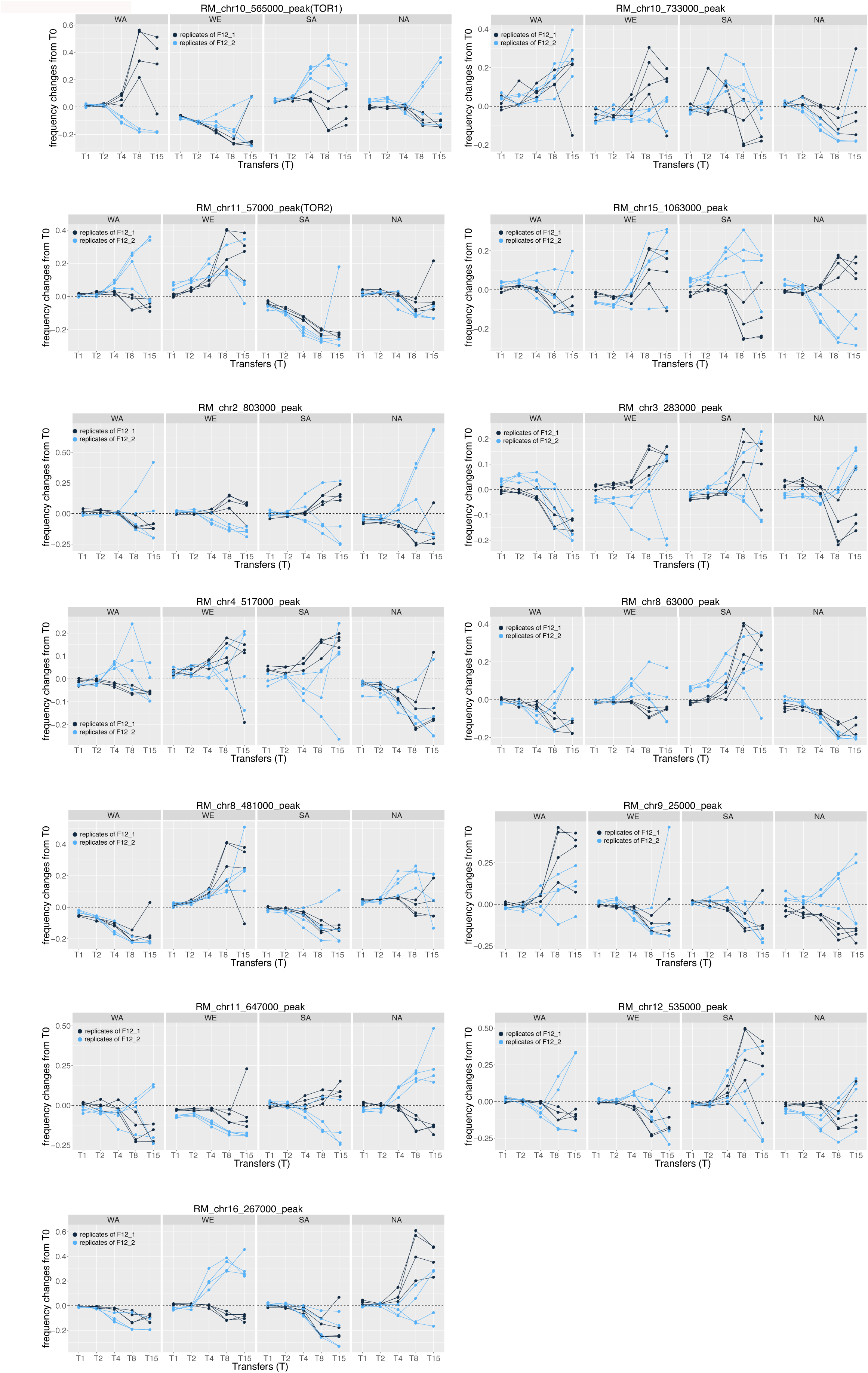
Allele frequencies dynamic at QTLs. Frequency changes of the parental alleles throughout the selection experiment with respect to their initial frequency at T0 in HU (A) and RM (B). Positive values indicate a frequency increases while negative values correspond to frequency decreases. Dots and lines in dark and light blue indicate replicates from F12_1 and F12_2 respectively.

**Figure S10.**
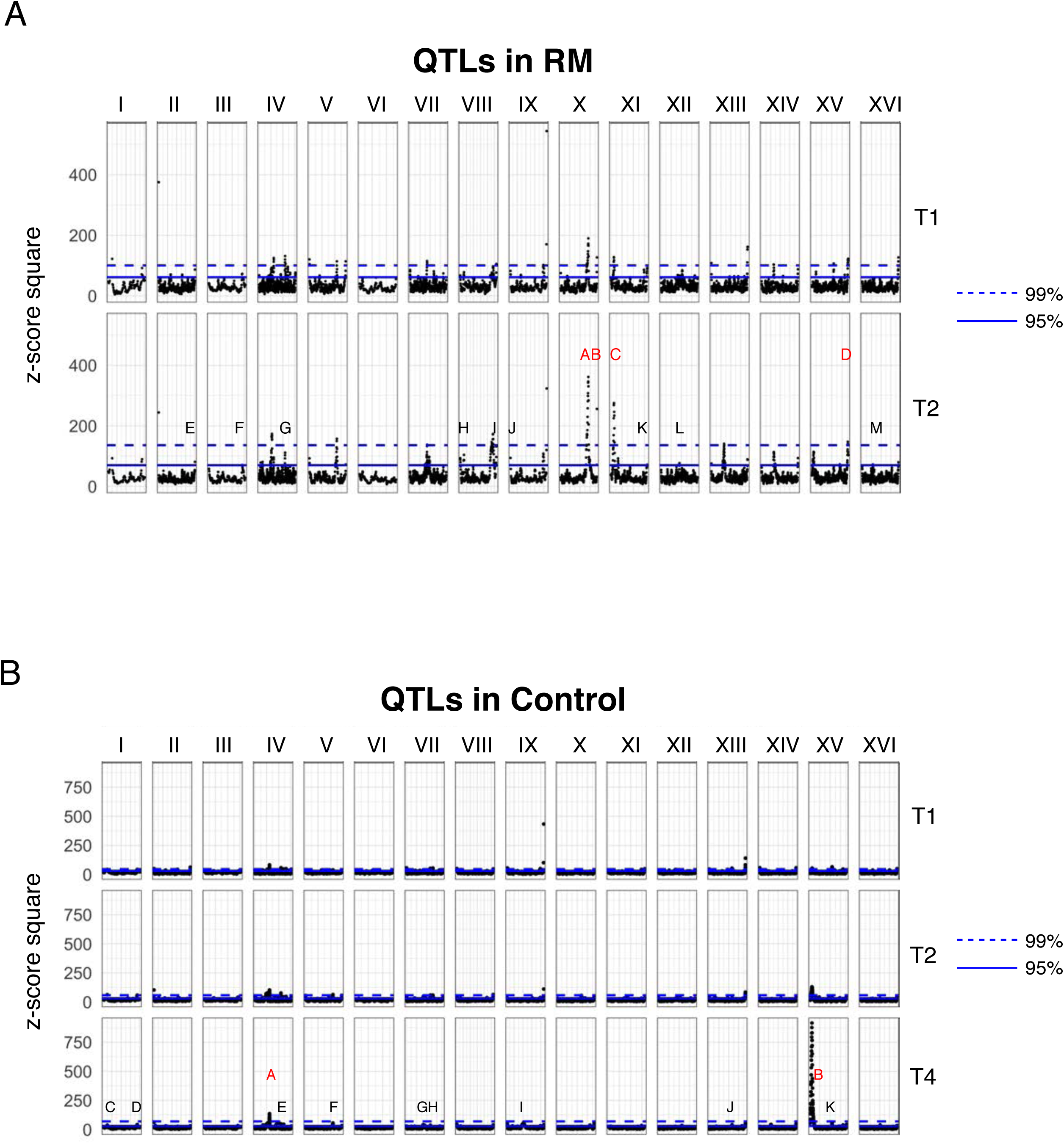
Identification of QTLs in RM and control conditions. We used allele frequency from T0 to T2 to identify QTLs for RM resistance (A) and T0 to T4 for drug-free condition (B). Dashed and solid lines indicate 99% and 95% quantile cut-off respectively. The red and black labels respectively indicate the QTLs passing 99% and 95% cut-off.

**Figure S11.**
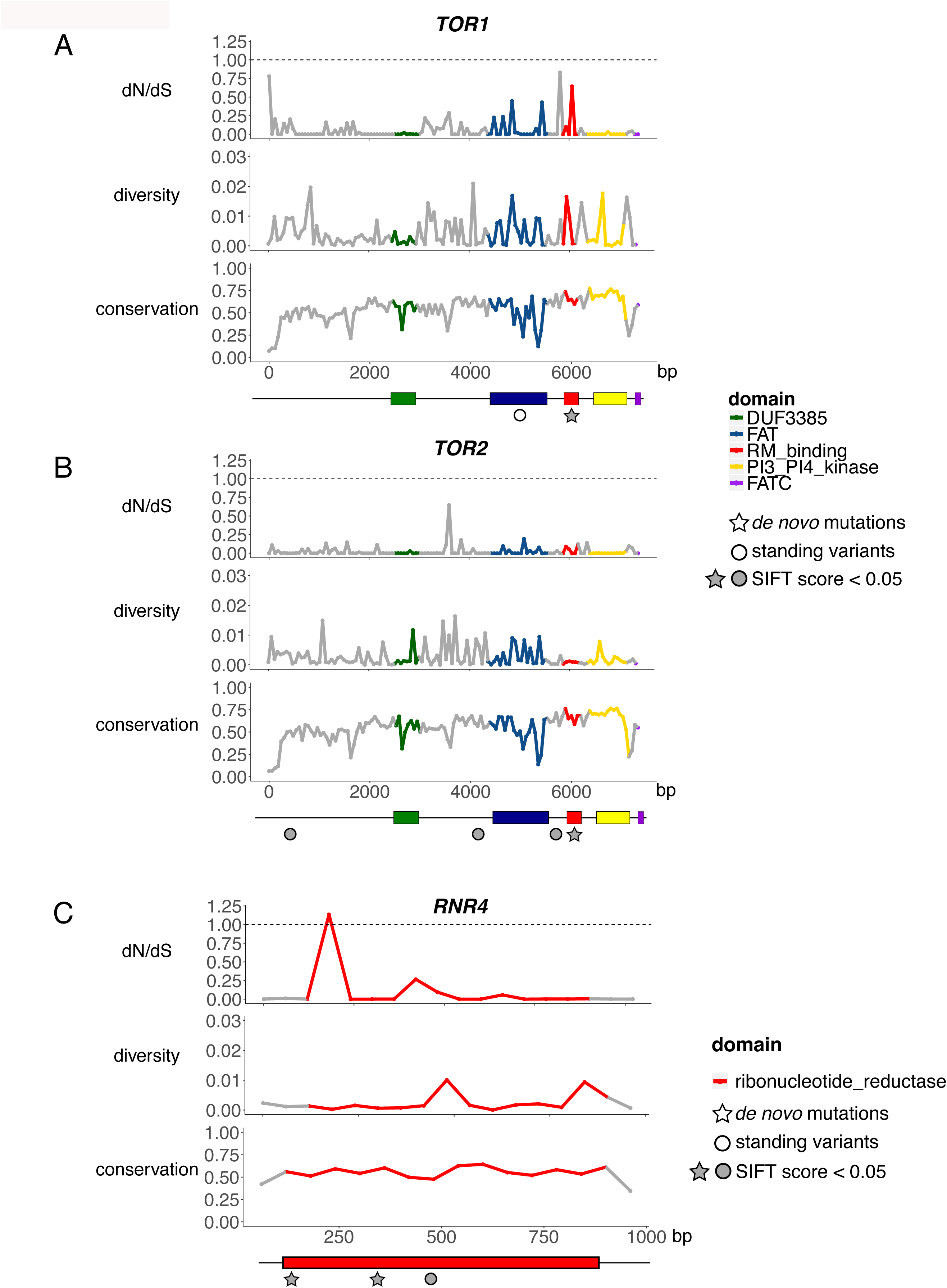
Sequence analysis of *TOR1, TOR2* and *RNR4*. The ratio of non-synonymous and synonymous substitutions (dN/dS), sequence diversity, and sequence conservation analysis of (A) *TOR1,* (B) *TOR2* and (C) *RNR4.* Functional domains are highlighted with different colors. The dashed line shows dN/dS = 1, indicating no selection (neutral). Values above 1 indicate positive selection and below 1 indicate purifying or stabilizing selection. All the plots are based on 60-bp window for each gene. dN/dS and diversity value is based on the sequences from the 1002 Yeast Genomes project. The conservation was calculated based on the multi-species sequence alignment compiled by SIFT for each tested polymorphic site (See Materials and Methods). Circle indicates the positions of standing variants and star indicates *de novo* mutations (Table 2). Grey color represents variants with significant SIFT score, indicating positions of high conservation.

**Figure S12.**
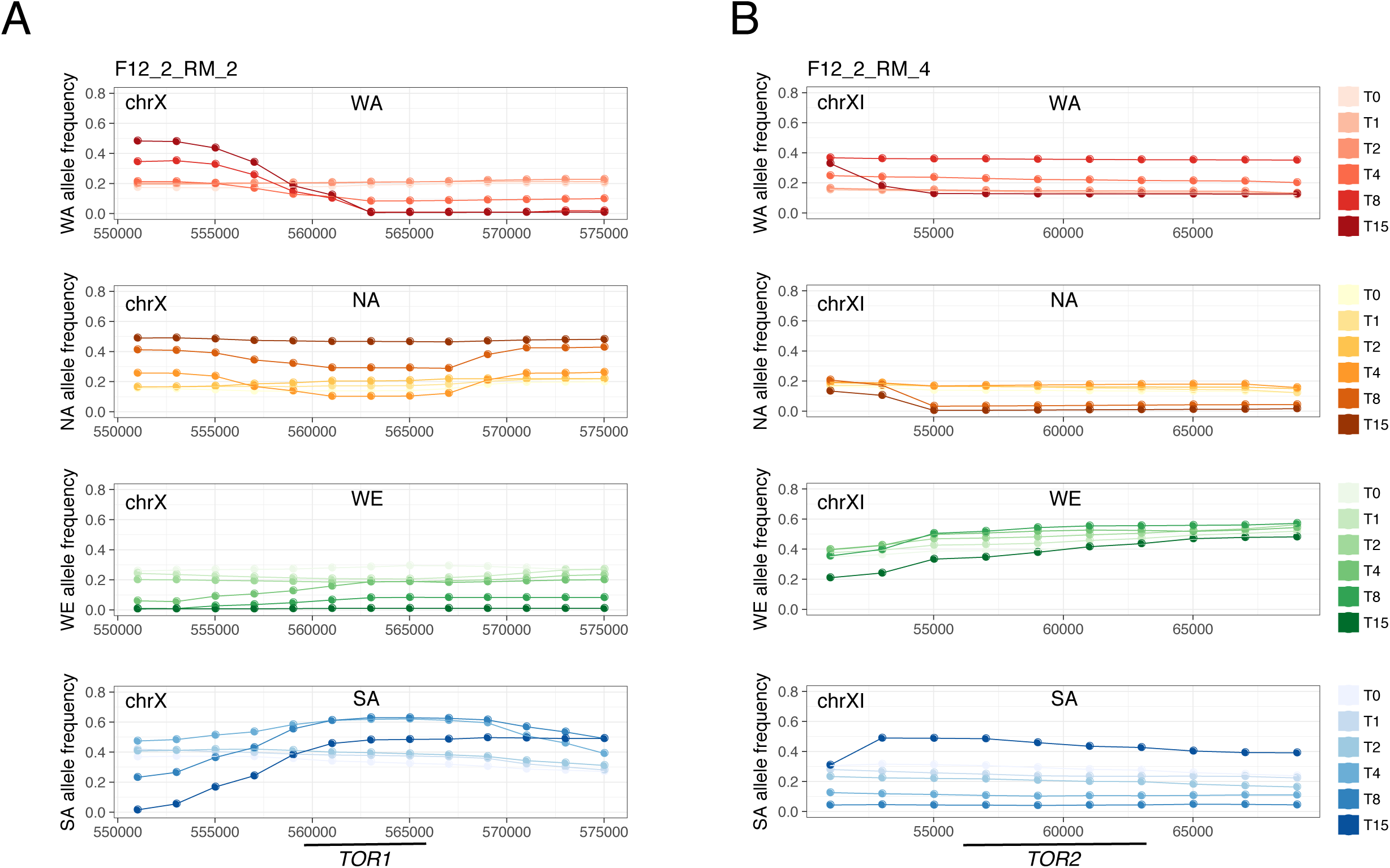
Local allele frequency changes in *TOR1* and *TOR2*. (A) Allele frequency changes of *TOR1* in population F12_2_RM_2. (B) Allele frequency changes of *TOR2* in population F12_2_RM_4. The panels show allele frequency changes of the four parental alleles (WA, NA, WE and SA from top to bottom). Early and late time points are indicated by colors from light to dark. The positions of *TOR1* and *TOR2* are indicated underneath the genomic coordinates reported in the x-axis.

**Figure S13.**
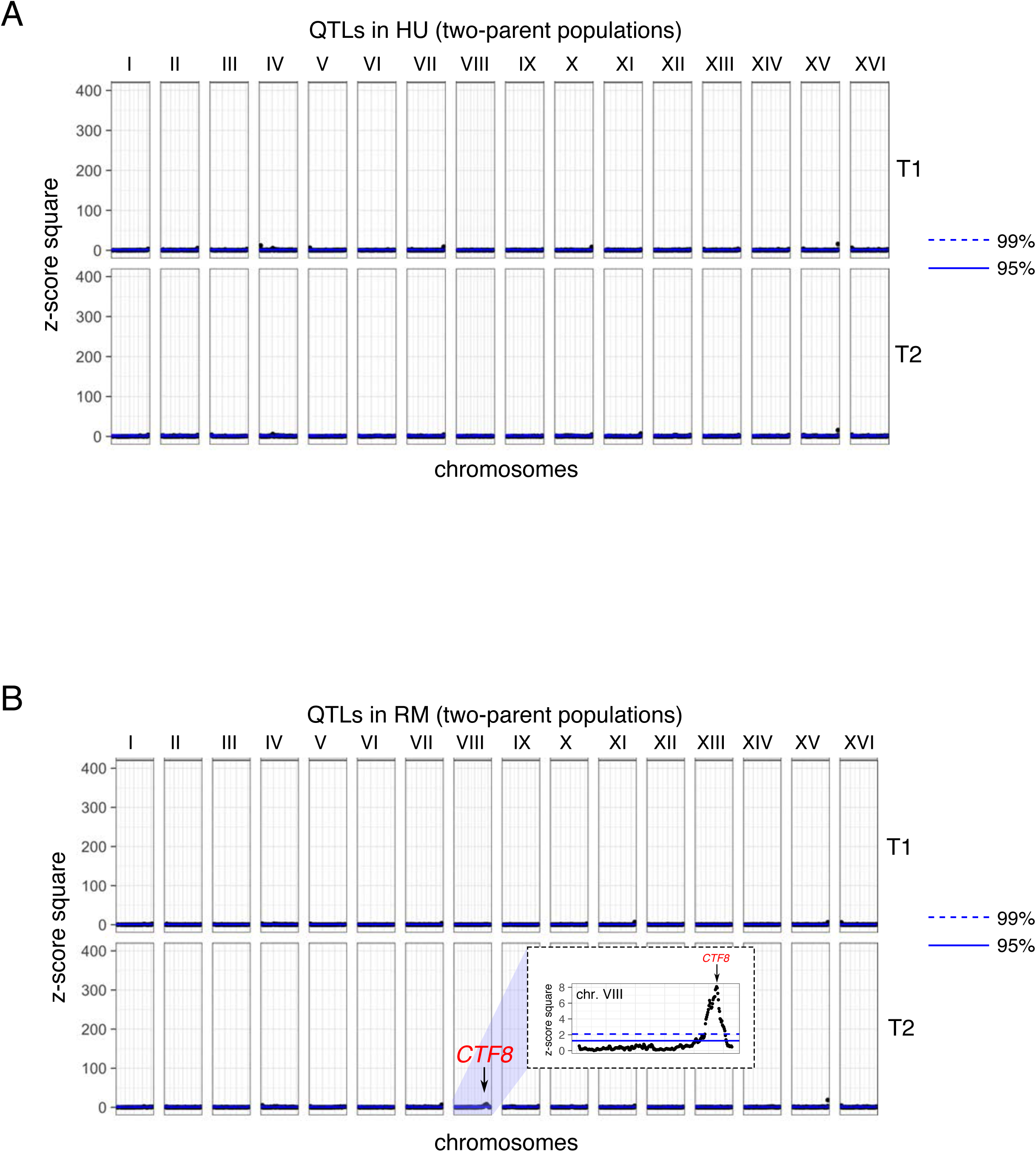
QTLs mapping in the two-parent populations. We applied the same QTL mapping approach used in this study to the two-parent populations dataset for HU (A) and RM (B). The z-score signals are very modest compared to the signals detected from the four-parent populations.

**Table S1.** Timeline of experimental evolution of isogenic, two-parent and four-parent populations.

| Transfer<br>(T) | Time lines (days) |  |  |  |  |  |
| --- | --- | --- | --- | --- | --- | --- |
|  | This study<br>(isogenic and four-parent populations) |  |  | Vazquez-Garcia, 2017<br>(two-parent populations) |  |  |
|  | HU | RM | Control | HU | RM | Control |
| **T0** | **0** | **0** | **0** | **0** | **0** | **0** |
| **T1** | **3** | **3** | **3** | **2** | **2** | **2** |
| **T2** | **6** | **6** | **6** | **4** | **4** | **4** |
| T3 | 9 | 8 | 8 | 6 | 6 | 6 |
| **T4** | **12** | **10** | **10** | **8** | **8** | **8** |
| T5 | 14 | 12 | 12 | 10 | 10 | 10 |
| T6 | 16 | 14 | 14 | 12 | 12 | 12 |
| T7 | 18 | 16 | 16 | 14 | 14 | 14 |
| **T8** | **20** | **18** | **18** | **16** | **16** | **16** |
| T9 | 22 | 20 | 20 | 18 | 18 | 18 |
| T10 | 24 | 22 | 22 | 20 | 20 | 20 |
| T11 | 26 | 24 | 24 | 22 | 22 | 22 |
| T12 | 28 | 26 | 26 | 24 | 24 | 24 |
| T13 | 30 | 28 | 28 | 26 | 26 | 26 |
| **T14** | **32** | 30 | 30 | 28 | 28 | 28 |
| **T15** |  | **32** | **32** | 30 | 30 | 30 |
| **T16** |  |  |  | **32** | **32** | **32** |

T0 to T16 indicate the number of serial transfers in the experimental evolution during the 32 days. We transferred cells of isogenic and four-parent cross populations every 2-3 days and made a total of 14 and 15 transfers in HU and RM, respectively. We transferred cells of the two-parent cross populations every 2 days and made a total of 16 transfers in both HU and RM (Vázquez-García et al., 2017). The labels in red indicate the time points when the populations were sequenced, which correspond to the initial population and the populations of the 1<sup>st</sup>, 2<sup>nd</sup>, 4<sup>th</sup>, 8<sup>th</sup> and the last transfer.

**Table S2.**
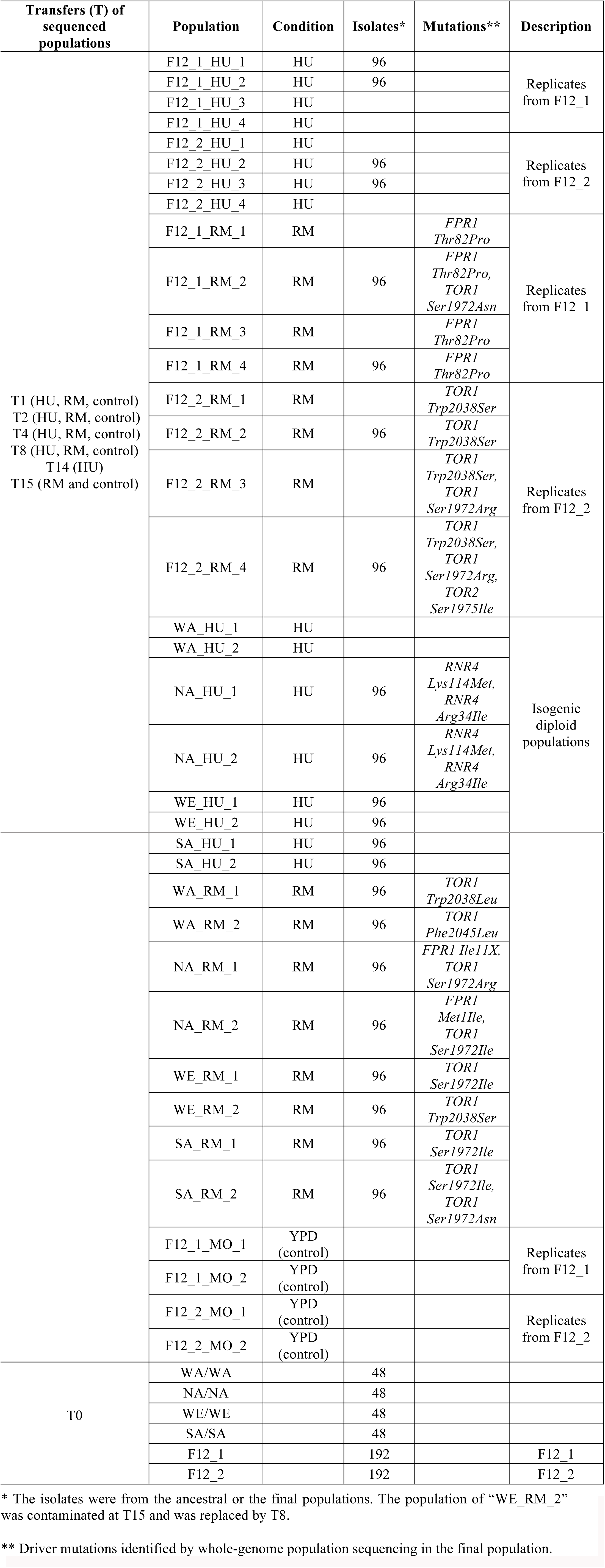
Summary of populations and isolates samples in this study.

| Transfers (T) of sequenced populations | Population | Condition | Isolates* | Mutations** | Description |
| --- | --- | --- | --- | --- | --- |
| T1 (HU, RM, control)<br>T2 (HU, RM, control)<br>T4 (HU, RM, control)<br>T8 (HU, RM, control)<br>T14 (HU)<br>T15 (RM and control) | F12_1_HU_1 | HU | 96 |  | Replicates from F12_1 |
|  | F12_1_HU_2 | HU | 96 |  |  |
|  | F12_1_HU_3 | HU |  |  |  |
|  | F12_1_HU_4 | HU |  |  |  |
|  | F12_2_HU_1 | HU |  |  | Replicates from F12_2 |
|  | F12_2_HU_2 | HU | 96 |  |  |
|  | F12_2_HU_3 | HU | 96 |  |  |
|  | F12_2_HU_4 | HU |  |  |  |
|  | F12_1_RM_1 | RM |  | *FPR1*<br>*Thr82Pro* | Replicates from F12_1 |
|  | F12_1_RM_2 | RM | 96 | *FPR1*<br>*Thr82Pro*,<br>*TOR1*<br>*Ser1972Asn* |  |
|  | F12_1_RM_3 | RM |  | *FPR1*<br>*Thr82Pro* |  |
|  | F12_1_RM_4 | RM | 96 | *FPR1*<br>*Thr82Pro* |  |
|  | F12_2_RM_1 | RM |  | *TOR1*<br>*Trp2038Ser* | Replicates from F12_2 |
|  | F12_2_RM_2 | RM | 96 | *TOR1*<br>*Trp2038Ser* |  |
|  | F12_2_RM_3 | RM |  | *TOR1*<br>*Trp2038Ser*,<br>*TOR1*<br>*Ser1972Arg* |  |
|  | F12_2_RM_4 | RM | 96 | *TOR1*<br>*Trp2038Ser*,<br>*TOR1*<br>*Ser1972Arg*,<br>*TOR2*<br>*Ser1975Ile* |  |
|  | WA_HU_1 | HU |  |  | Isogenic diploid populations |
|  | WA_HU_2 | HU |  |  |  |
|  | NA_HU_1 | HU | 96 | *RNR4*<br>*Lys114Met*,<br>*RNR4*<br>*Arg34Ile* |  |
|  | NA_HU_2 | HU | 96 | *RNR4*<br>*Lys114Met*,<br>*RNR4*<br>*Arg34Ile* |  |
|  | WE_HU_1 | HU | 96 |  |  |
|  | WE_HU_2 | HU | 96 |  |  |
|  | SA_HU_1 | HU | 96 |  |  |
|  | SA_HU_2 | HU | 96 |  |  |
|  | WA_RM_1 | RM | 96 | *TOR1*<br>*Trp2038Leu* |  |
|  | WA_RM_2 | RM | 96 | *TOR1*<br>*Phe2045Leu* |  |
|  | NA_RM_1 | RM | 96 | *FPR1 Ile11X,*<br>*TOR1*<br>*Ser1972Arg* |  |
|  | NA_RM_2 | RM | 96 | *FPR1*<br>*Met1Ile,*<br>*TOR1*<br>*Ser1972Ile* |  |
|  | WE_RM_1 | RM | 96 | *TOR1*<br>*Ser1972Ile* |  |
|  | WE_RM_2 | RM | 96 | *TOR1*<br>*Trp2038Ser* |  |
|  | SA_RM_1 | RM | 96 | *TOR1*<br>*Ser1972Ile* |  |
|  | SA_RM_2 | RM | 96 | *TOR1*<br>*Ser1972Ile,*<br>*TOR1*<br>*Ser1972Asn* |  |
|  | F12_1_MO_1 | YPD<br>(control) |  |  | Replicates<br>from F12_1 |
|  | F12_1_MO_2 | YPD<br>(control) |  |  |  |
|  | F12_2_MO_1 | YPD<br>(control) |  |  | Replicates<br>from F12_2 |
|  | F12_2_MO_2 | YPD<br>(control) |  |  |  |
| T0 | WA/WA |  | 48 |  |  |
|  | NA/NA |  | 48 |  |  |
|  | WE/WE |  | 48 |  |  |
|  | SA/SA |  | 48 |  |  |
|  | F12_1 |  | 192 |  | F12_1 |
|  | F12_2 |  | 192 |  | F12_2 |

\* The isolates were from the ancestral or the final populations. The population of “WE\_RM\_2” was contaminated at T15 and was replaced by T8.
\*\* Driver mutations identified by whole-genome population sequencing in the final population.

Table S3. Sequencing depth (median) of all the samples

See separated file

Table S4. List of genotyped clones

See separated file

**Table S5.**
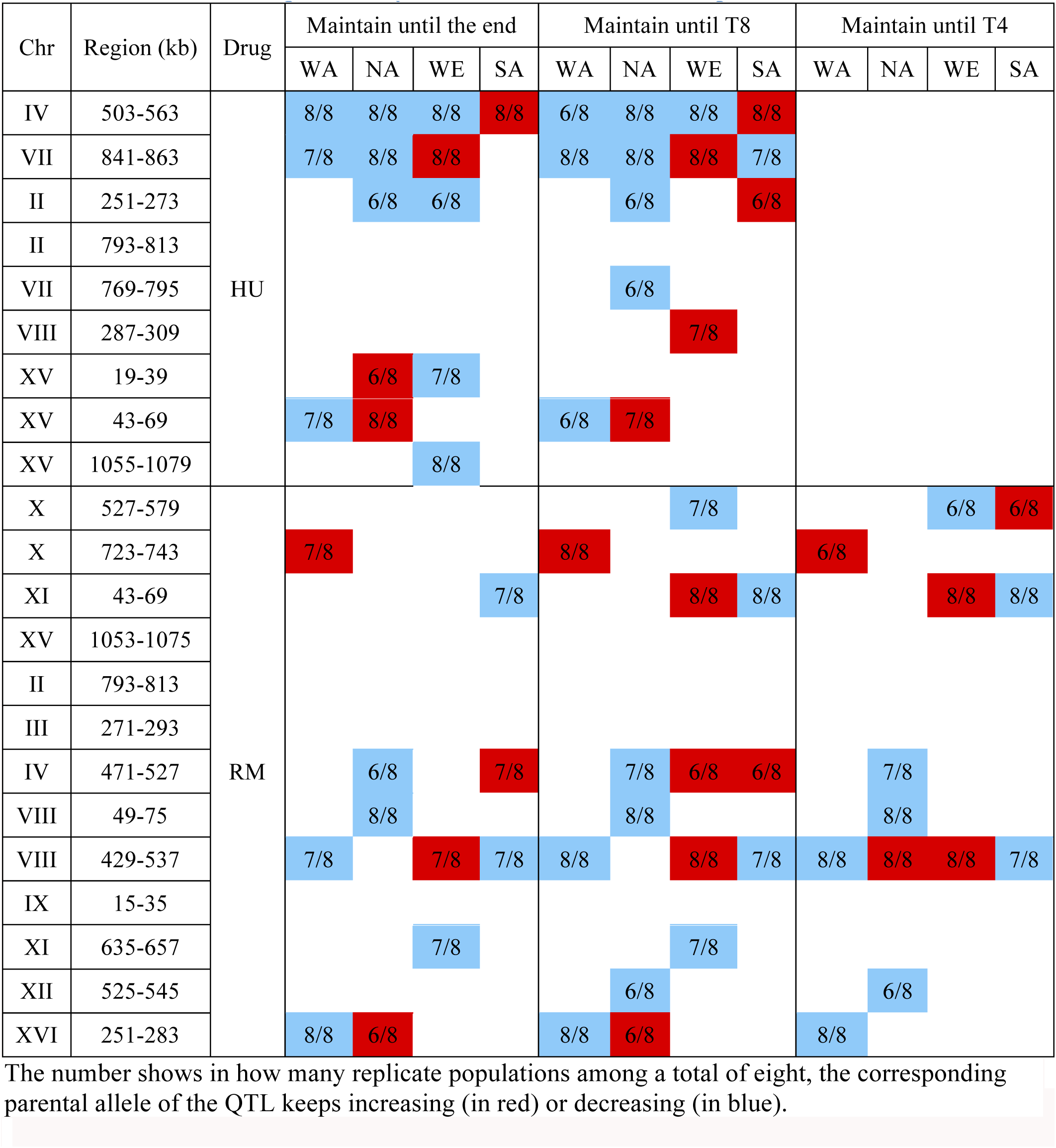
QTL allele frequencies dynamic at late selection time points.

Table S6 Functional variants identified in 1002 Yeast Genomes project

See separated file

**Table S7.** Tetrad viability analysis of *TOR2* hemizygous deletions.

| Background | Medium | Tetrads dissected | Distribution of tetrad types |  |  |  |  | Spore viability |
| --- | --- | --- | --- | --- | --- | --- | --- | --- |
|  |  |  | 4-sv | 3-sv | 2-sv | 1-sv | 0-sv |  |
| SA/WE *tor2Δ* | SC | 96 | 0.03 | 0.26 | 0.61 | 0.06 | 0.03 | 0.55 |
| SA *tor2Δ*/WE | SC | 82 | 0.02 | 0.39 | 0.41 | 0.13 | 0.04 | 0.56 |
| SA/WE | SC | 35 | 0.49 | 0.40 | 0.09 | 0.03 | 0.00 | 0.84 |
| SA/WE *tor2Δ* | YPD | 24 | 0.00 | 0.04 | 0.67 | 0.25 | 0.04 | 0.43 |
| SA *tor2Δ*/WE | YPD | 22 | 0.00 | 0.00 | 0.86 | 0.14 | 0.00 | 0.47 |
| SA/WE | YPD | 470 | 0.53 | 0.18 | 0.15 | 0.04 | 0.10 | 0.75 |

3 The value displayed in each “Distribution of tetrad types” column is the frequency of tetrads 4 containing four viable spores (4-sv), three viable spores (3-sv), two viable spores (2-sv), one 5 viable spore (1-sv) and no viable spores (0-sv). The far right column shows the overall spore 6 viability of each strain.

**Table S8.** Number of inferred modifiers for *TOR2* dispensability.

| Background | Tetrads dissected | Parental ditype (2:2) | Nonparental ditype (4:0) | Tetratype (3:1) | Single gene *p* value | Three gene *p* value | Wild type *p* value |
| --- | --- | --- | --- | --- | --- | --- | --- |
| SA/WE *tor2*Δ | 96 | 59 | 6 | 31 | 7.14E-31 | 0 | 5.56E-77 |
| SA *tor2*Δ/WE | 82 | 34 | 5 | 43 | 4.97E-09 | 0 | 3.39E-30 |
| SA/WE | 35 | 3 | 17 | 15 | 2.59E-06 | 2.40E-13 | - |

2 Based on the tetrad segregation pattern, a chi-squared statistic ( $\chi^2$ ) was used to test three separate 3 hypothesis: (1) a single unlinked modifier explains the inheritance patterns (1:1:4 ratio expected); 4 (2) three unlinked modifiers explain the inheritance patterns; and (3) many loci make the 5 inheritance patterns indistinguishable from empirically observed background. In all cases, a *p* 6 value was calculated for the $\chi^2$ statistic using `chisq.test` in R. See (Dowell et al., 2010) for detail 7 to calculate the *p* value.

**Table S9.**
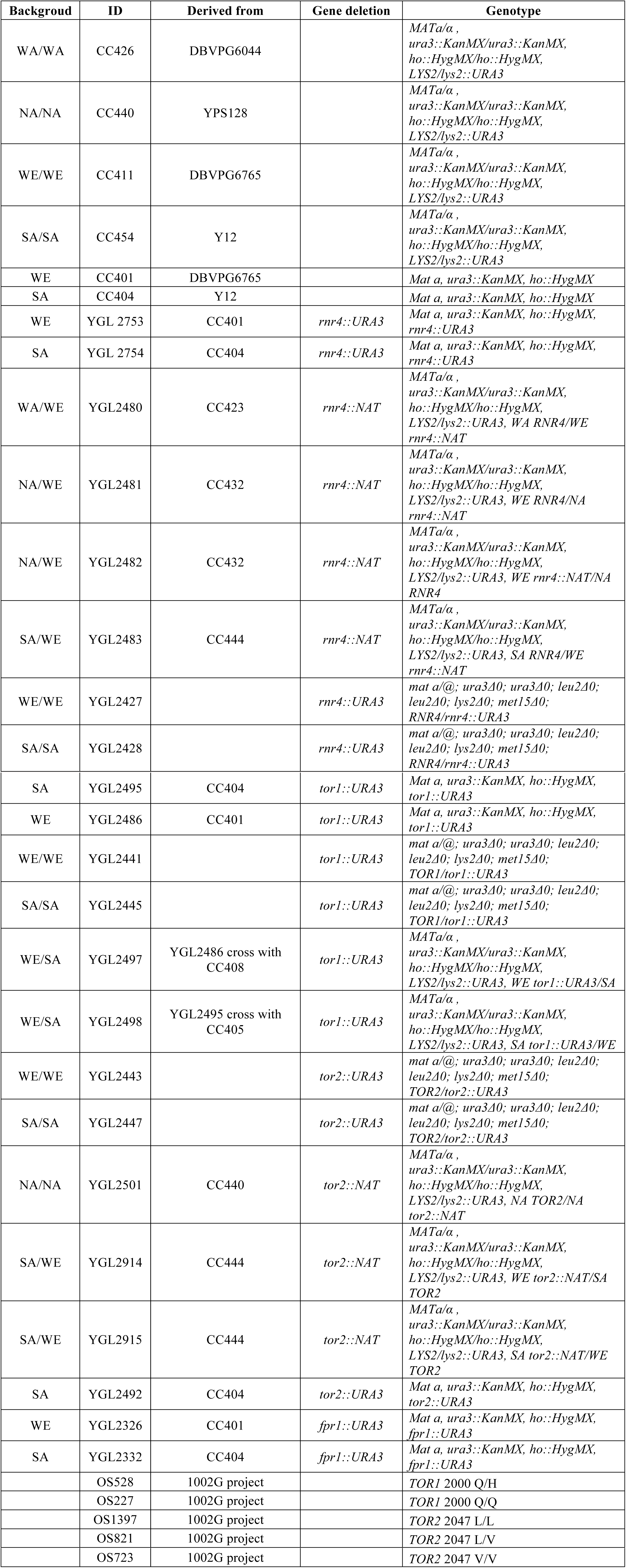
Strains used in this study.

**Table S10.**
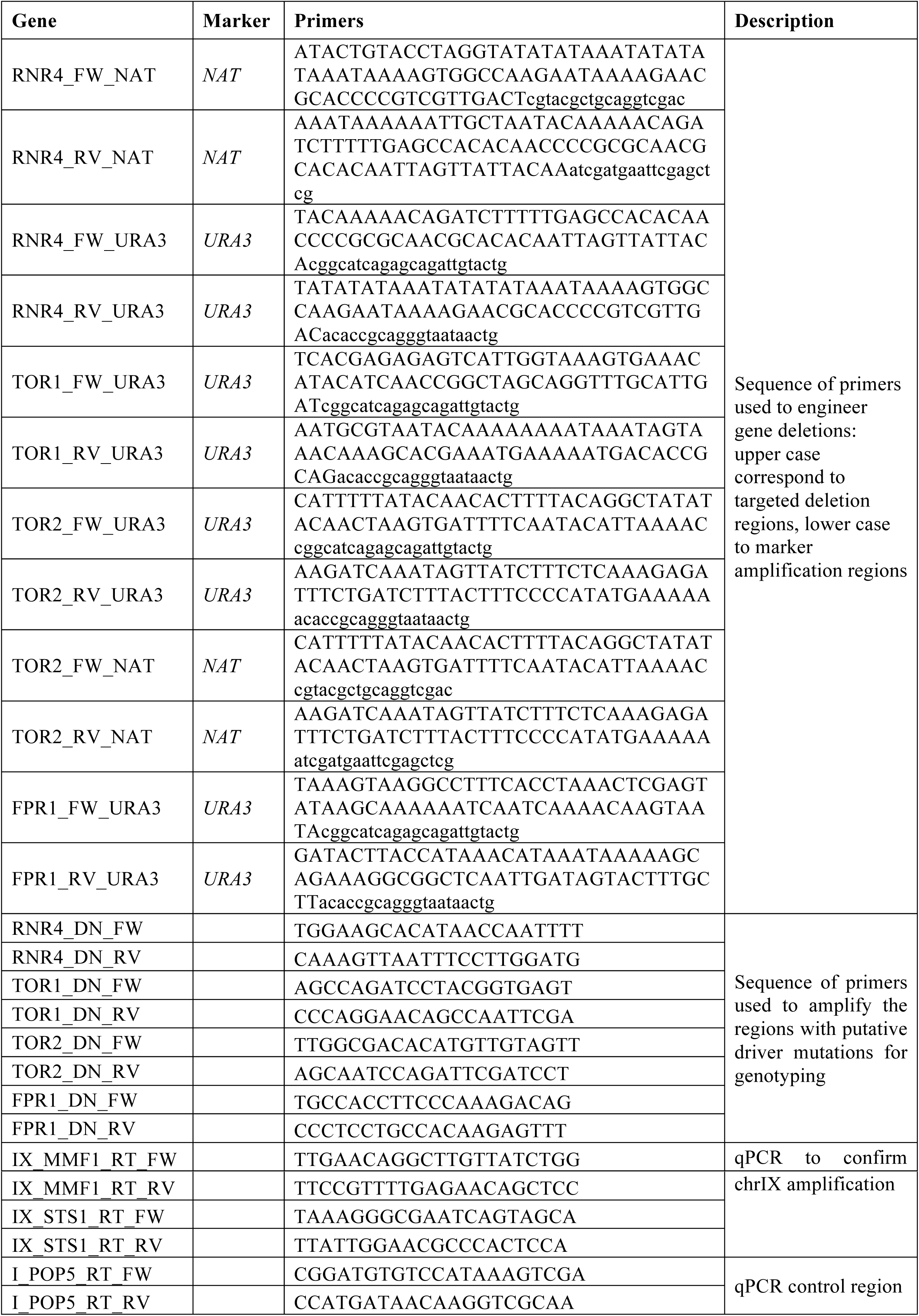
Primers used in this study.

